# Reclassification of Pterulaceae Corner (Basidiomycota: Agaricales) introducing the ant-associated genus *Myrmecopterula* gen. nov., *Phaeopterula* Henn. and the corticioid Radulomycetaceae fam. nov.

**DOI:** 10.1101/718809

**Authors:** Caio A. Leal-Dutra, Gareth W. Griffith, Maria Alice Neves, David J. McLaughlin, Esther G. McLaughlin, Lina A. Clasen, Bryn T. M. Dentinger

## Abstract

Pterulaceae was formally proposed to group six coralloid and dimitic genera [*Actiniceps* (=*Dimorphocystis*), *Allantula, Deflexula, Parapterulicium, Pterula* and *Pterulicium*]. Recent molecular studies have shown that some of the characters currently used in Pterulaceae Corner do not distinguish the genera. *Actiniceps* and *Parapterulicium* have been removed and a few other resupinate genera were added to the family. However, none of these studies intended to investigate the relationship between Pterulaceae genera. In this study, we generated 278 sequences from both newly collected and fungarium samples. Phylogenetic analyses support by morphological data allowed a reclassification of Pterulaceae where we propose the introduction of *Myrmecopterula* gen. nov. and Radulomycetaceae fam. nov., the reintroduction of *Phaeopterula*, the synonymisation of *Deflexula* in *Pterulicium* and 51 new combinations. *Pterula* is rendered polyphyletic requiring a reclassification; thus, it is split into *Pterula, Myrmecopterula* gen. nov., *Pterulicium* and *Phaeopterula. Deflexula* is recovered as paraphyletic alongside several *Pterula* species and *Pterulicium*, and is sunk into the latter genus. *Phaeopterula* is reintroduced to accommodate species with darker basidiomes. The neotropical *Myrmecopterula* gen. nov. forms a distinct clade adjacent to *Pterula*, and most members of this clade are associated with active or inactive attine ant nests. The resupinate genera *Coronicium* and *Merulicium* are recovered in a strongly supported clade close to *Pterulicium*. The other resupinate genera previously included in Pterulaceae, and which form basidiomes lacking cystidia and with monomitic hyphal structure (*Radulomyces, Radulotubus* and *Aphanobasidium*), are reclassified into Radulomycetaceae fam. nov. *Allantula* is still an enigmatic piece in this puzzle known only from the type specimen that requires molecular investigation. A key for the genera of Pterulaceae and Radulomycetaceae fam. nov. is provided here.

## INTRODUCTION

The history of Pterulaceae Corner begins with the hesitant proposal of the genus *Pterula* Fr. (hereinafter abbreviated as Pt.) in the 1820s and 1830s by Elias Magnus Fries (Fries 1821, 1825, 1830). The typification of this genus was addressed by Lloyd (1919) and this was followed by discussion between M.S. Doty, M.A. Donk and D.P. Rogers (Doty 1948; Donk 1949; Rogers 1949, 1950). Ultimately Corner (1952c) provided a thorough discussion of the timeline of Fries’ decisions, which was later confirmed with further clarification by M.A. Donk (Donk 1954; Donk 1963).

The number of species in *Pterula* grew during the late 19th and early 20th centuries, with J.H. Léveille, N.T Patouillard, P. C. Hennings, P. A. Saccardo, C. G. Lloyd, C. L. Spegazzini and M. J. Berkeley being the most active in the naming of taxonomic novelties of *Pterula* in this period (Corner 1950, 1970). Lloyd (1919) devoted an entire chapter to discuss the taxonomy of the genus. However, the major contribution to the genus was made by E. J. H. Corner who added at least 45 new taxa (Corner 1950, 1952b, 1970, 1966, 1967). Corner (1950) created the *Pteruloid* series in Clavariaceae Chevall. to group, besides *Pterula*, other genera with coralloid basidiome and dimitic hyphal system. The *Pteruloid* series was raised by Donk (1964) to Pteruloideae, a sub-family of Clavariaceae. Pterulaceae was formally proposed by Corner (1970) including the genera from the original Pteruloideae: Allantula Corner, *Deflexula* Corner, *Dimorphocystis* Corner (=*Actiniceps* Berk. & Broome), *Parapterulicium* Corner, *Pterula* and *Pterulicium* Corner (hereinafter abbreviated as Pm.) (Corner 1950, 1952a, b, 1970) (Fig. 1).

**Fig. 1.**
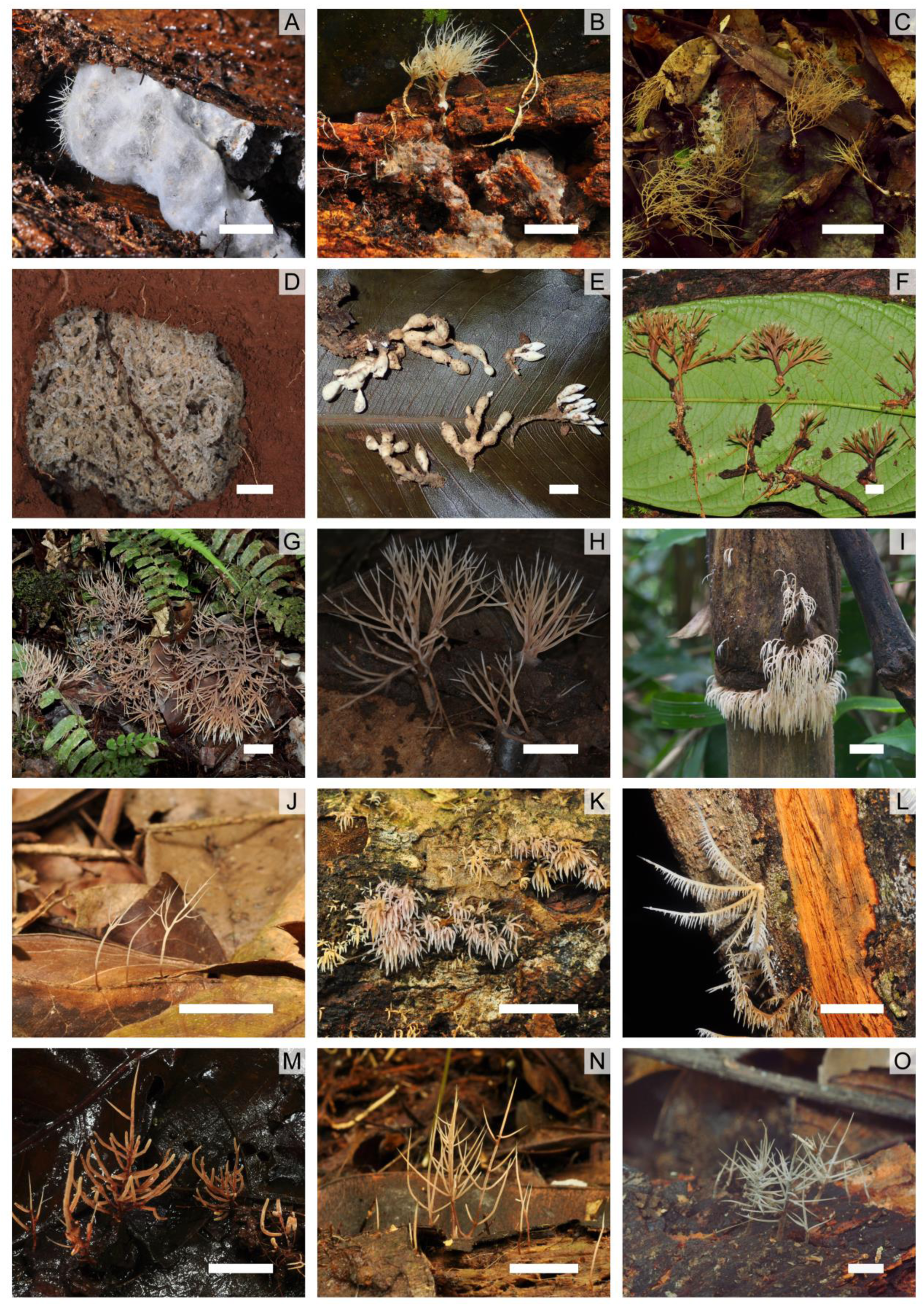
Diversity of coralloid genera of Pterulaceae. A-F: *Myrmecopterula* (A: *Apterostigma sp*. nest with *M. velohortorum* with *Myrmecopterula sp*. SAPV1 growing atop of the garden veil; B, C, F: *Myrmecopterula sp*. D: *Apterostigma sp*. nest with *M. nudihortorum*; E: *M. moniliformis*). G-H: *Pterula* (G: *P. cf. loretensis*; H: *P. cf. verticillata*). I-L: *Pterulicium* (I: *P. secundirameum*; J: *P. aff fluminensis*; K: *P. lilaceobrunneum*. L: *P. sprucei*). M-O: *Phaeopterula* (M: *Phaeopterula sp*.; N: *P. stipata*; O: *P. juruensis*). Close observation on photos B and C reveal the basidiomes growing from granular substrate resembling substrate of ants’ fungus garden. Photos D, E and G kindly provided by Ted Schultz, Susanne Sourell and Michael Wherley respectively. Scale bars: 1 cm.

Following Corner’s reclassifications, the major changes in Pterulaceae have resulted from molecular phylogenetic analyses. *Actiniceps* Berk. & Broome was shown within *Agaricales* to be distantly related to Pterulaceae and *Parapterulicium* was removed to *Russulales* (Dentinger and McLaughlin 2006; Leal-Dutra et al. 2018). Four resupinate genera were transferred to Pterulaceae: *Aphanobasidium* Jülich, Coronicium J. Erikss. & Ryvarden, *Merulicium* J. Erikss. & Ryvarden, and *Radulomyces* M.P. Christ. (Larsson 2007; Larsson et al. 2004) and, finally, the new poroid genus *Radulotubus* Y.C. Dai, S.H. He & C.L. Zhao was proposed in the family (Zhao et al. 2016) (Fig. 2).

**Fig. 2.**
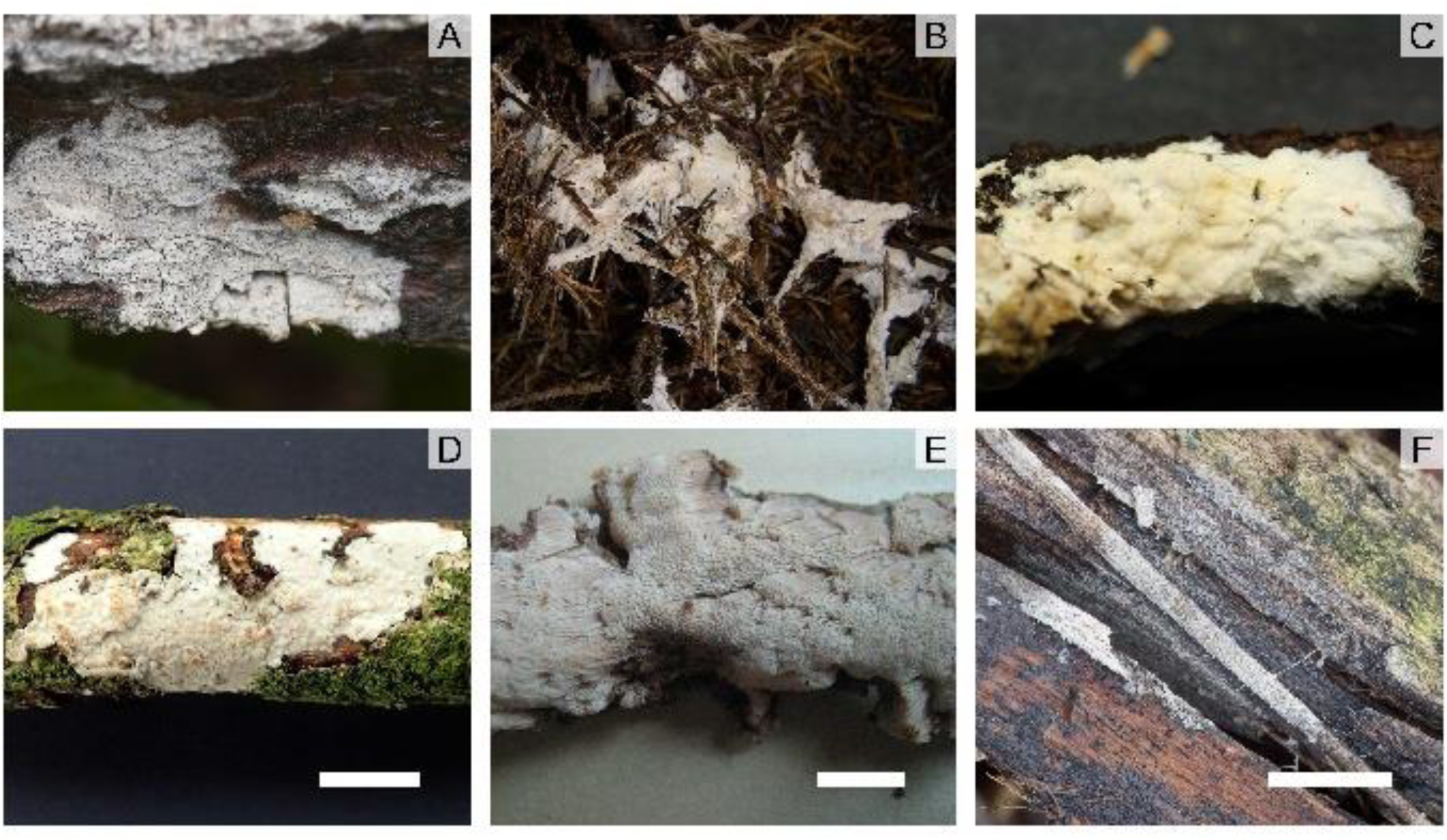
Corticioid genera of Pterulaceae (A-C) and Radulomycetaceae (D-F). A: *Coronicium alboglaucum*. B-C: *Merulicium fusisporum*. D: *Radulomyces confluens*. E: *Radulotubus resupinatus*. F: *Aphanobasidium cf. pseudotsugae*. Photos kindly provided by L. Zíbarová (A and F), S. Blaser (B and C), D.J. Harries (D) and C.L. Zhao (E). Scale bars: 1 cm.

The ecological roles of Pterulaceae are not well understood, most being classified from superficial observations as saprotrophs, growing on wood or leaf litter, with wood decay potentially being the ancestral state. Whilst many species are found inhabiting soil or litter, two species are reported to associate with living plants, namely *Pterula cf. tenuissima*, endophytic in asymptomatic leaves of *Magnolia grandiflora*, and *Pterulicium xylogenum*, causal agent of culm rot disease of bamboo (Munkacsi et al. 2004; Villesen et al. 2004; Harsh et al. 2005) and possibly also a pathogen of sugarcane [see Corner (1952b)].

Pterulaceae has attracted more attention recently following the discovery of two distinct symbionts of fungus-farming ants in the genus *Apterostigma* Mayr being included in several phylogenetic and ecological studies (Matheny et al. 2006; Hibbett 2007; Dentinger et al. 2009; Binder et al. 2010; Leal-Dutra 2015). Despite the absence (hitherto) of any teleomorph, phylogenetic analyses placed both species [*Pterula nudihortorum* Dentinger and *Pterula velohortorum* Dentinger, previously known as G2 and G4 (Dentinger 2014)] in a strongly supported clade within Pterulaceae (Munkacsi et al. 2004; Villesen et al. 2004).

Whilst these earlier phylogenetic studies did not focus on resolving evolutionary relationships of the genera, they did demonstrate that the coralloid genera of Pterulaceae are clearly polyphyletic. Amongst the morphological characters previously used to separate the genera, but now known to be phylogenetically unreliable, is the orientation of basidiome growth that differentiates *Pterula* from *Deflexula* and the presence of a corticioid patch at the base of the basidiome in *Pterulicium* (Corner 1950, 1952a, 1970). Therefore, the reclassification of Pterulaceae is required to restore the monophyly of the genera.

We aimed to clarify the phylogenetic relationships of the various genera within Pterulaceae through collection of new samples during fieldwork campaigns in Brazil and additionally sampling of fungarium specimens. This has yielded sequence data from many specimens not included in previously phylogenetic analyses, permitting a comprehensive reappraisal of the phylogeny of Pterulaceae. Here we present a proposal for a new classification based on the phylogeny inferred from three nuclear loci (nrITS, nrLSU and RPB2), including representatives of all genera currently accepted in Pterulaceae except *Allantula*. Despite several attempts for recollecting *Allantula* in its type locality, the monotypic genus is still only known from the type specimen collected by Corner (1952a).

## METHODS

### Collections and morphological observations

Several field campaigns between 2011-2017 have obtained new specimens from >15 locations in nine states across Brazil (Amazonas, Espírito Santo, Minas Gerais, Pará, Paraíba, Paraná, Rio de Janeiro, Rio Grande do Sul and Santa Catarina). The samples were dried in a low-heat food dehydrator and deposited at Universidade Federal de Santa Catarina, Universidade Federal do Oeste do Pará, Instituto Nacional de Pesquisas da Amazônia and Royal Botanic Gardens, Kew [FLOR, HSTM, INPA, K and RB, respectively]; herbarium acronyms follow Index Herbariorum (Thiers continuously updated)]. Morphological identification and taxonomy of Pterulaceae are treated sensu Corner. Microscopic observations followed the methods described in Leal-Dutra (2015) and Leal-Dutra et al. (2018).

### DNA extraction, amplification, cloning and sequencing

DNA was extracted from dried basidiomes or freeze-dried culture by first grinding with liquid nitrogen and then lysis in CTAB buffer (100 mM Tris-HCl pH 8.0, 1.4 M NaCl, 20 mM EDTA, 2% CTAB), clean-up with chloroform:isoamyl alcohol (24:1), wash with isopropanol (0.6 vol.) and a final wash with 70% ethanol. Partial sequences of the nrITS, nrLSU and RPB2 were amplified by PCR using the primer pairs listed on Table 1 and following the cycling conditions in the original publications. PCR products were purified using 2 U of Exonuclease I (Thermo Fisher Scientific) and 0.2 U FastAP Thermosensitive Alkaline Phosphatase (Thermo Fisher Scientific) per 1 µl of PCR product, incubated at 37°C for 15 min, followed by heat inactivation at 85 °C for 15 min. The samples were then sent for Sanger sequencing at the IBERS Translational Genomics Facility (Aberystwyth University) or Jodrell Laboratory (Royal Botanic Gardens, Kew). The same PCR primers were used for sequencing; additional primers were used to sequence the nrLSU and RPB2 (Table 1).

**Table 1.**
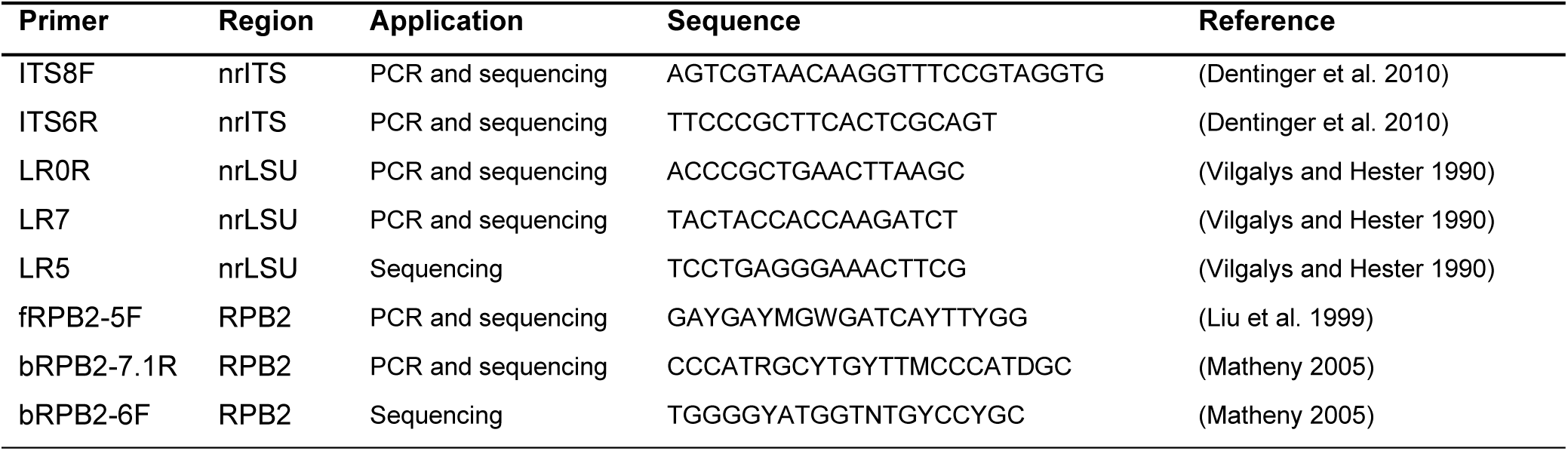
Primers used in this study for PCR and sequencing.

Chromatograms were checked and sequences assembled and edited using GENEIOUS 10.0.2 (Kearse et al. 2012). Samples presenting indels were cloned using pGEM-T Easy Vector Systems (Promega) into Subcloning Efficiency DH5α Competent Cells (Invitrogen). Up to five clones from each sample were amplified and sequenced as above. For each sample clone sequences were aligned to generate one or more consensus sequences and polymorphisms were replaced by respective IUPAC code for ambiguous nucleotide; in cases where indels were found, two different sequences were saved [see Leal-Dutra et al. (2018)].

Moreover, 27 sequences of nrITS (4), nrLSU (10) and RPB2 (13) were mined from 13 previously assembled and unpublished genomes using NCBI BLAST+ package v2.7.1 (Camacho et al. 2009). Two sequences of each Pterulaceae genus were used as query and the best hit based on the combination of e-value and bit score was selected; the same hit should usually appear for all query sequences. In one case (sample KM190547), more than one optimal hit was found; the subject sequences were compared for occurrence of indels and treated as virtual clones (VC). These sequences are included in the dataset (Table 2). The sequences generated in this study have been submitted to GenBank (Table 2).

**Table 2:** Details of new sequences generated in this study used in the tree of Fig. 3.

| **Taxon (former genus in brackets)** | **DNA sample ID** | **Herbarium voucher** | **Country** | **Region** | **ITS** | **LSU** | **RPB2** |
| --- | --- | --- | --- | --- | --- | --- | --- |
| *Coronicium alboglaucum* | K15 | K(M) 170129 | UK | England | MK953245 | – | – |
| *Coronicium gemmiferum* | K13 | K(M) 133847 | UK | England | MK953246 | – | – |
| *Coronicium gemmiferum* | K14 | K(M) 68853 | UK | England | MK953247 | MK953403 | – |
| *Merulicium fusisporum* | K16 | K(M) 45181 | UK | England | MK953248 | – | – |
| *Myrmecopterula (Pterula) moniliformis* | F92 Consensus 1 | INPA 280127 | Brazil | Amazonas | MK953251 | MK953405 | MK944362 |
| *Myrmecopterula (Pterula) moniliformis* | M39 | FLOR 56397 | Brazil | Paraíba | MK953253 | MK953406 | – |
| *Myrmecopterula (Pterula) moniliformis* | MCA | not deposited | – | – | MK953239 | MK953392 | MK944363 |
| *Myrmecopterula (Pterula) nudihortorum* | F144 Consensus 1 | not deposited | Brazil | Amazonas | MK953257 | MK953393 | MK944364 |
| *Myrmecopterula (Pterula) nudihortorum* | KM190547_VC1 | K(M) 190547 | Panama | – | MK953240 | MK953394 | MK944365 |
| *Myrmecopterula (Pterula) sp.* | F103 | HSTM-Fungos 9931 | Brazil | Pará | MK953260 | MK953407 | MK944325 |
| *Myrmecopterula (Pterula) sp.* | F138 | FLOR 63724 | Brazil | Paraná | MK953262 | MK953409 | – |
| *Myrmecopterula (Pterula) sp.* | F40 | FLOR 63725 | Brazil | Paraná | MK953264 | MK953410 | MK944327 |
| *Myrmecopterula (Pterula) sp.* | F82 Consensus 1 | not deposited | Brazil | Amazonas | MK953269 | MK953412 | MK944366 |
| *Myrmecopterula (Pterula) sp.* | F94 | HSTM-Fungos 9928 | Brazil | Pará | MK953274 | MK953414 | MK944329 |
| *Myrmecopterula (Pterula) sp.* | F99 | HSTM-Fungos 9930 | Brazil | Pará | MK953276 | MK953415 | MK944330 |
| *Myrmecopterula (Pterula) sp.* | M111 | FLOR 56451 | Brazil | Santa Catarina | MK953277 | – | – |
| *Myrmecopterula (Pterula) sp.* | M40 Consensus 1 | FLOR 56398; K(M) 205347 | Brazil | Paraíba | MK953280 | MK953416 | MK944367 |
| *Myrmecopterula (Pterula) sp.* | M69 | FLOR 56418 | Brazil | Rio Grande do Sul | MK953281 | MK953395 | MK944368 |
| *Myrmecopterula (Pterula) velohortorum* | F114 | not deposited | Brazil | Espírito Santo | MK953282 | MK953396 | MK944369 |
| *Myrmecopterula (Pterula) velohortorum* | F117 | not deposited | Brazil | Santa Catarina | MK953283 | – | – |
| *Myrmecopterula (Pterula) velohortorum* | F135 | not deposited | Brazil | Pará | MK953285 | – | – |
| *Myrmecopterula (Pterula) velohortorum* | F136 | not deposited | Brazil | Pará | MK953286 | – | – |
| *Myrmecopterula (Pterula) velohortorum* | F137 | not deposited | Brazil | Pará | MK953287 | – | – |
| *Myrmecopterula (Pterula) velohortorum* | F140 Clone 1 | not deposited | Brazil | Amazonas | MK953288 | – | – |
| *Myrmecopterula (Pterula) velohortorum* | F152 | not deposited | Brazil | Santa Catarina | MK953290 | – | – |
| *Myrmecopterula (Pterula) velohortorum* | KM190546 | K(M) 190546 | Panama | – | MK953242 | MK953397 | MK944370 |
| *Myrmecopterula (Pterula) velohortorum* | RC12 Consensus 1 | not deposited | Brazil | Amazonas | MK953291 | – | – |
| *Phaeopterula (Pterula) anomala* | KM38182 | K(M) 38182 | Cameroon | – | MK953295 | – | – |
| *Phaeopterula (Pterula) cf. juruensis* | F45 Consensus 1 | FLOR 63732 | Brazil | Paraná | MK953296 | MK953417 | MK944331 |
| *Phaeopterula (Pterula) cf. juruensis* | F79 Consensus 1 | FLOR 63717 | Brazil | Paraná | MK953299 | MK953418 | – |
| *Phaeopterula (Pterula) cf. stipata* | F66 Consensus 1 | HSTM-Fungos 9938 | Brazil | Pará | MK953301 | – | – |
| *Phaeopterula (Pterula) cf. stipata* | F98 Consensus 1 | HSTM-Fungos 9929 | Brazil | Pará | MK953302 | – | – |
| *Phaeopterula (Pterula) cf. taxiformis* | M4 | FLOR 56367 | Brazil | Santa Catarina | MK953303 | MK953419 | – |
| *Phaeopterula (Pterula) juruensis* | F41 | FLOR 63728 | Brazil | Paraná | MK953304 | MK953420 | MK944332 |
| *Phaeopterula (Pterula) juruensis* | M21 | FLOR 56381 | Brazil | Minas Gerais | MK953305 | – | – |
| *Phaeopterula (Pterula) juruensis* | M36 | FLOR 56396 | Brazil | Santa Catarina | MK953307 | MK953422 | – |
| *Phaeopterula (Pterula) sp.* | F63 Consensus 1 | HSTM-Fungos 9935 | Brazil | Pará | MK953316 | MK953425 | MK944335 |
| *Phaeopterula (Pterula) sp.* | F78 Clone 1 | FLOR 63716 | Brazil | Paraná | MK953321 | MK953428 | MK944338 |
| *Phaeopterula (Pterula) sp.* | KM135954 | K(M) 135954 | Belize | – | MK953326 | – | – |
| *Phaeopterula (Pterula) sp.* | KM137475 | K(M) 137475 | Puerto Rico | – | MK953327 | – | – |
| *Phaeopterula (Pterula) stipata* | M15 Consensus 1 | FLOR 56375 | Brazil | Minas Gerais | MK953330 | MK953431 | – |
| *Phaeopterula sp. (Allantula?)* | F7 Consensus 1 | HSTM-Fungos 9944 | Brazil | Pará | MK953331 | MK953432 | MK944339 |
| *Pterula cf. plumosa* | KM167176 | K(M) 167176 | Ethiopia | – | MK953333 | – | – |
| *Pterula cf. loretensis* | RLC273 | K(M) 205553 | Ecuador | Imbabura | MK953334 | MK953398 | MK944371 |
| *Pterula multifida* | KM195746 | K(M) 195746 | UK | England | MK953335 | MK953399 | MK944372 |
| *Pterula sp.* | F42 | FLOR 63729 | Brazil | Paraná | MK953336 | MK953433 | – |
| *Pterula sp.* | F48 | FLOR 63735 | Brazil | Paraná | – | MK953434 | – |
| *Pterula sp.* | M112 Consensus 1 | FLOR 56452 | Brazil | Santa Catarina | MK953337 | MK953435 | MK944340 |
| *Pterula sp.* | M153 | FLOR 57849 | Brazil | Santa Catarina | MK953339 | MK953436 | MK944341 |
| *Pterula sp.* | M54 | FLOR 56407 | Brazil | Santa Catarina | MK953341 | MK953438 | – |
| *Pterula sp.* | M71 Consensus 1 | FLOR 56424 | Brazil | Santa Catarina | MK953342 | MK953439 | MK944342 |
| *Pterula sp.* | KM141379 | K(M) 141379 | Puerto Rico | – | MK953344 | – | – |
| *Pterula sp.* | KM167221 | K(M) 167221 | Australia | Queensland | MK953345 | – | – |
| *Pterula subulata* | KM145950 | K(M) 145950 | Italy | – | MK953346 | – | – |
| *Pterula subulata* | KM167186 | K(M) 167186 | Sweden | – | MK953347 | – | – |
| *Pterula verticillata* | KM27119 | K(M) 27119 | Brunei | – | MK953348 | – | – |
| *Pterulicium (Deflexula) fasciculare* | KM167225 | K(M) 167225 | Australia | – | MK953349 | – | – |
| *Pterulicium (Deflexula) fasciculare* | KM167227 | K(M) 167227 | Malaysia | – | MK953350 | – | – |
| *Pterulicium (Deflexula) lilaceobrunneum* | M117 | FLOR 56455 | Brazil | Rio de Janeiro | MK953351 | MK953440 | MK944343 |
| *Pterulicium (Deflexula) secundirameum* | BZL44 | RB 575791 | Brazil | Rio de Janeiro | MK953353 | MK953400 | MK944373 |
| *Pterulicium (Deflexula) secundirameum* | M50 | FLOR 56403 | Brazil | Santa Catarina | MK953354 | MK953442 | MK944344 |
| *Pterulicium (Deflexula) sp.* | KM167228 | K(M) 167228 | Malaysia | – | MK953357 | – | – |
| *Pterulicium (Deflexula) sp.* | KM167233 | K(M) 167233 | Sierra Leone | – | MK953358 | – | – |
| *Pterulicium (Deflexula) sprucei* | F68 | HSTM-Fungos 9940 | Brazil | Pará | MK953361 | MK953447 | MK944349 |
| *Pterulicium (Deflexula) subsimplex* | KM160100 | K(M) 160100 | Ecuador | – | – | MK953449 | – |
| *Pterulicium (Deflexula) subsimplex* | M33 | FLOR 56393 | Brazil | Santa Catarina | MK953363 | MK953450 | MK944351 |
| *Pterulicium (Pterula) brunneosetosum* | M35 Consensus 1 | FLOR 56395 | Brazil | Santa Catarina | MK953366 | MK953452 | MK944353 |
| *Pterulicium (Pterula) caricispendulae* | KM155784 | K(M) 155784 | UK | England | MK953367 | – | – |
| *Pterulicium (Pterula) sp.* | F20 | INPA 280129 | Brazil | Amazonas | MK953370 | MK953454 | – |
| *Pterulicium (Pterula) sp.* | F21 | INPA 280132 | Brazil | Amazonas | MK953371 | MK953455 | MK944355 |
| *Pterulicium (Pterula) sp.* | F26 | not deposited | Brazil | Espírito Santo | MK953372 | MK953456 | MK944356 |
| *Pterulicium (Pterula) sp.* | F30 | not deposited | Brazil | Espírito Santo | MK953373 | MK953457 | MK944357 |
| *Pterulicium (Pterula) sp.* | F57 | HSTM-Fungos 9925 | Brazil | Pará | MK953376 | MK953460 | MK944359 |
| *Pterulicium (Pterula) sp.* | F76 Consensus 1 | HSTM-Fungos 9950 | Brazil | Pará | MK953382 | MK953461 | MK944360 |
| *Pterulicium (Pterula) sp.* | M1 | FLOR 56364 | Brazil | Santa Catarina | MK953383 | MK953462 | MK944361 |
| *Pterulicium (Pterula) sp.* | M6 | FLOR 56369 | Brazil | Santa Catarina | MK953384 | MK953463 | – |
| *Pterulicium (Pterulicium) xylogenum* | KM167222 | K(M) 167222 | Bangladesh | – | MK953387 | – | – |
| *Aphanobasidium pseudotsugae* | K6 | K(M) 170662 | UK | England | MK953243 | MK953402 | – |
| *Aphanobasidium pseudotsugae* | K7 | K(M) 180787 | UK | Scotland | MK953244 | – | – |
| *Radulomyces confluens* | KM167249 | K(M) 167249 | Brazil | – | MK953388 | – | – |
| *Radulomyces confluens* | KM167250 | K(M) 167250 | Argentina | – | MK953389 | – | – |
| *Radulomyces confluens* | KM181613 | K(M) 181613 | UK | England | MK953390 | MK953401 | MK944374 |
| *Radulomyces copelandii* | M150 | K(M) 173275 | USA | – | MK953391 | MK953465 | – |

### Phylogenetic analyses

A preliminary maximum-likelihood (ML) analysis was conducted with the sequences generated in this study alongside GenBank sequences to find the best outgroup for Pterulaceae based on previous studies (Dentinger et al. 2016; Zhao et al. 2016; Matheny et al. 2006; Larsson 2007) and to assess the similarities between the cloned sequences (Additional file 1; Additional file 2).

A reduced version of the previous dataset with only one sequence from each cloned sample was created after removing nearly identical sequences with no phylogenetic resolution. The final dataset was comprised of 119 sequences, including 32 sequences from GenBank and using four sequences of Stephanospora Pat. as outgroups, and was divided into five partitions for further analyses: ITS1, 5.8S, ITS2, LSU and RPB2. Each partition was aligned separately with MAFFT v7.311 (Katoh and Standley 2013) using the E-INS-i algorithm for ITS1 and ITS2, and L-INS-i for 5.8S, LSU and RPB2. The alignments were examined and corrected manually in AliView v1.5 (Larsson 2014) and trimmed to remove uneven ends. Following the simple indel coding (Simmons and Ochoterena 2000), a morphological matrix were constructed using SeqState (Müller 2005) where indels were coded as binary characters. The nucleotide alignments were then trimmed with trimAl v1.4.rev22 (Capella-Gutiérrez et al. 2009) with the option - gappyout to remove unaligned regions.

Maximum-likelihood tree reconstruction was performed with IQ-TREE v1.6.7.1 (Nguyen et al. 2015). The best-fit evolutionary models and partitioning scheme for this analysis were estimated by the built-in ModelFinder (option -m MF+MERGE) allowing the partitions to share the same set of branch lengths but with their own evolution rate (-spp option) (Chernomor et al. 2016; Kalyaanamoorthy et al. 2017). Branch support was assessed with 1000 replicates of ultrafast bootstrapping (UF-Boot) (Hoang et al. 2018) and allowing resampling partitions and then sites within these partitions to reduce the likelihood of false positives on branch support (option - bspec GENESITE).

Bayesian Inference (BI) was implemented using MRBAYES v3.2 (Ronquist et al. 2012) with two independent runs, each one with four chains and starting from random trees. The best-fit evolutionary models and partitioning scheme for these analyses were estimated as for the ML analysis but restricting the search to models implemented on MRBAYES (options - m TESTMERGEONLY -mset mrbayes). Chains were run for 107 generations with tree sampling every 1000 generations. The burn-in was set to 25% and the remaining trees were used to calculate a 50% majority consensus tree and Bayesian Posterior Probability (BPP). The convergence of the runs was assessed on TRACER v1.7 (Rambaut et al. 2018) to ensure the potential scale reduction factors (PSRF) neared 1.0 and the effective sample size values (ESS) were sufficiently large (>200). Nodes with BPP ≥0.95 and/or UFBoot ≥95 were considered strongly supported. Alignment and phylogenetic trees are deposited in Treebase (ID: 24428).

## RESULTS

From this section, all taxa are treated by the nomenclatural treatment proposed in this study.

### Field data

Fieldwork resulted in the discovery of approximately 100 new specimens, now placed within Pterulaceae (Table 2). Axenic culture isolation was also possible from several of these specimens.

### Phylogenetic analyses

A total of 278 sequences from 123 samples were generated in this study: 153 nrITS, 74 nrLSU and 51 RPB2; 61 from cloning and 40 from genome mining. The final alignment consisted of 113 sequences with 2737 characters and 1050 parsimony-informative sites. The BI analysis converged both runs as indicated by the effective sample sizes (ESS) of all parameters above 2800 and the potential scale reduction factors (PSRF) equal 1.000 for all the parameters according to the 95% HPD Interval.

The new classification proposed in this study (Fig. 3), highlights six main clades containing nine genera are highlighted: Radulomycetaceae (containing *Aphanobasidium, Radulotubus* and *Radulomyces*), *Phaeopterula* (hereinafter abbreviated as Ph.; previously *Pterula* spp.), *Coronicium* superclade (grouping *Merulicium* and *Coronicium*), *Pterulicium* (previously *Pterulicium, Pterula* spp. and *Deflexula* spp,), *Pterula* and *Myrmecopterula* (*Myrmecopterula* gen. nov., previously *Pterula* spp.).

**Fig. 3.**
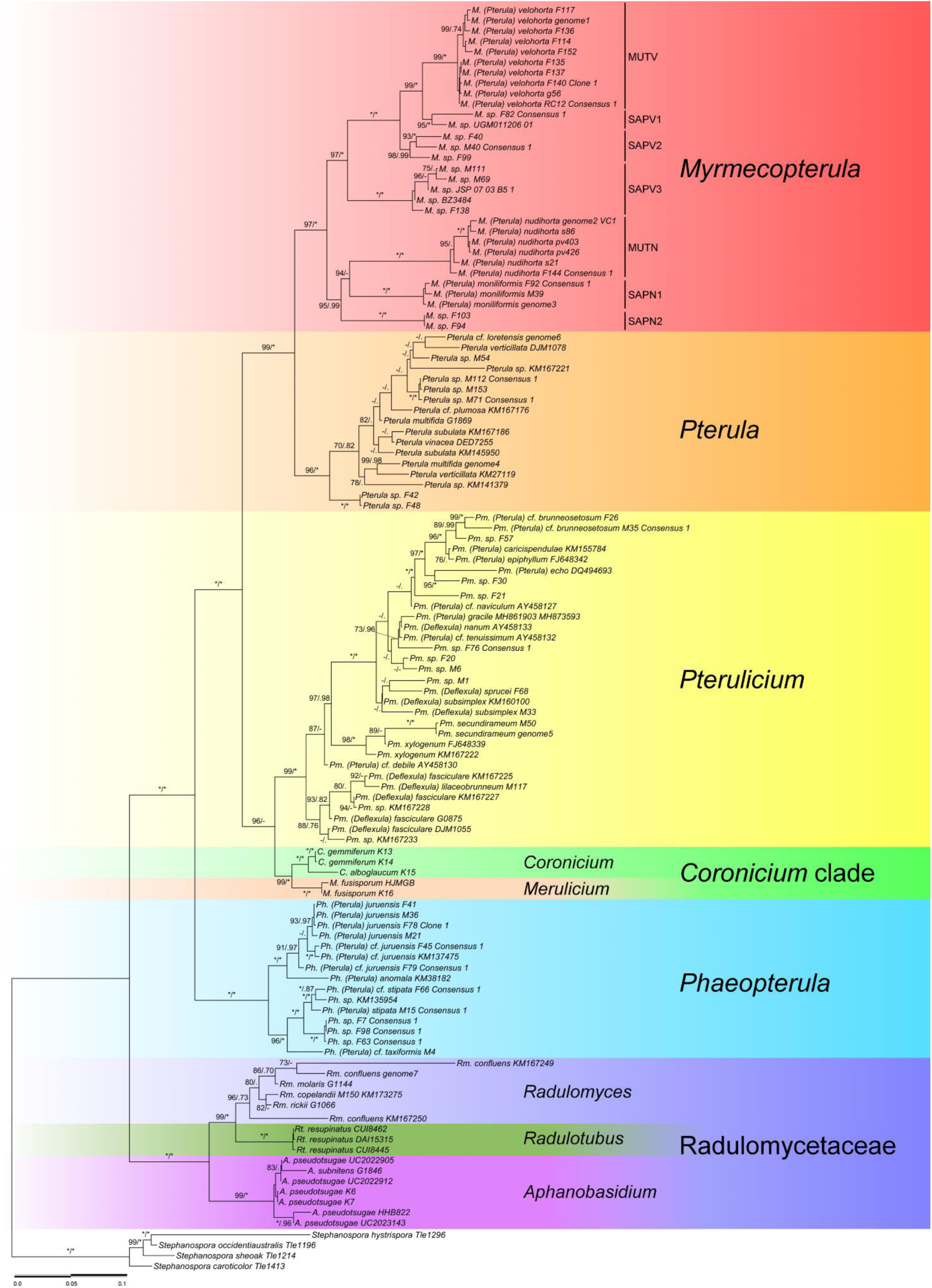
Maximum-likelihood tree of Pterulaceae and Radulomycetaceae. Support values on the branches are UF-Boot/BPP and shown only for UFBoot≥70 and BPP≥0.70 and branch length ≥ 0.003 substitutions per site. Asterisks (*) represent maximum UFBoot/BPP values, dashes (-) represent values below the cut-off threshold (70%), and dots (.) represent ML clades that was not recovered in BI tree. Details for the complete tree can be found in Additional file 2 and TreeBase (ID: 24428). Scale bar: nucleotide substitutions per site.

### Radulomycetaceae (UFBoot=99; BPP=1)

This clade groups with strong support three of the five resupinate genera recognized in Pterulaceae, namely *Aphanobasidium* (UF-Boot=100; BPP=1), Radulotubus (UF-Boot=100; BPP=1) and Radulomyces (UF-Boot=100; BPP=0.86). The placement of Aphanobasidium and *Radulomyces* into Pterulaceae was previously shown by phylogenetic reconstructions of corticioid taxa (Larsson et al. 2004; Larsson 2007). *Radulotubus* was proposed by Zhao et al. (2016) as sister clade of

*Radulomyces* to accommodate one species bearing poroid hymenophore. In our analyses, *Radulotubus* was recovered in the same position as in the original publication. This is the only poroid species within Pterulaceae.

No members of the three genera within this superclade are pteruloid (i.e. coralloid basidiomes with dimitic hyphal system) in their morphology and consequently we propose the creation of Radulomycetaceae fam. nov. to accommodate them, as discussed in greater detail below. The current sister clade to Pterulaceae in our analyses is Stephanosporaceae Oberw. & E.Horak, from which members of the Radulomycetaceae clade are clearly distinct phylogenetically and morphologically.

### Phaeopterula (UFBoot=100; BPP=1)

*Phaeopterula* received maximum support in both analyses. It includes *Pterula stipata* Corner, *Pterula anomala* P. Roberts, *Pterula juruensis* (Henn.) Corner and other species which all have dark brown basidiomes. This clade is the first coralloid lineage to diverge within Pterulaceae. As these species render *Pterula* paraphyletic, a reclassification is needed. The generic name *Phaeopterula* (Henn.) Sacc. & D. Sacc was originally proposed as a subgenus of *Pterula* to accommodate *Ph. hirsuta* Henn. and *Ph. juruensis* (Hennings 1904; Hennings 1900). We propose its reintroduction to distinguish these brown-pigmented taxa from *Pterula* s.s.

### Coronicium superclade (UFBoot=98; BPP=1)

This clade groups the remaining two resupinate genera of Pterulaceae, the monospecific *Merulicium* and *Coronicium* (UFBoot=100; BPP=1). Both genera form resupinate basidiomes but differ in the hyphal system present (dimitic in *Merulicium, monomitic* in *Coronicium*). Some *Pterulicium* species also show transitions in their morphology to a resupinate state. Corner (1950) showed that *Pm. xylogenum* (Berk. & Broome) Corner could form monomitic corticioid patches independent of the coralloid state and even in its absence, thus appearing to be truly corticioid. Furthermore, experimental studies on *Pm. echo* D.J. McLaughlin & E.G. McLaughlin show a dimitic, resupinate, fertile corticioid phase both on agar and when cultured on cocoa twigs (McLaughlin and McLaughlin 1980; McLaughlin et al. 1978; McLaughlin and McLaughlin 1972). Despite the morphological distinctiveness from the rest of Pterulaceae, there is a trend in the morphology and strong phylogeny support for the placement of the *Coronicium* superclade among *Pterula/Myrmecopterula* and *Pterulicium* clades within Pterulaceae.

### Pterulicium (UFBoot=99; BPP=1)

Two type species, *Pterulicium xylogenum* and *Deflexula fascicularis*, are nested within this clade alongside several species currently assigned to *Pterula* but which all have simple basidiomes (unbranched or limited branching). The *Pterula* species are interspersed with some *Deflexula*, rendering both genera polyphyletic. *Pterulicium xylogenum* forms a well-supported subclade with *Pterula secundiramea* (Lév.) Corner (=Pm. palmicola Corner). *Deflexula fascicularis* forms a subclade with other *Deflexula* species that share globose spores, an unusual feature within Pterulaceae, most of which form ellipsoid to subamygdaliform spores.

### Pterula (UFBoot=100; BPP=1)

This clade groups the true *Pterula* spp. that are represented by very bushy coralloid basidiomes, usually robust and taller than those of *Pterulicium*, stipe concolorous with hymenophore and lacking a cottony subiculum. *Pterula* has a mainly pantropical and pan-subtropical distribution, with occurrence reported to all continents except Antarctica (Corner 1970).

### Myrmecopterula (UFBoot=97; BPP=1)

This sister clade of *Pterula* represents the newly proposed genus (see below). It groups the two species cultivated by attine ants in the *Apterostigma pilosum* group with *M. moniliformis* and several unidentified free-living species. The species in this clade are only known from the Neotropics. *Myrmecopterula* is divided in seven subclades (Fig. 3) representing the two mutualists (MUT 1-2), three closely related to *M. velohortorum* (SAPV 1-3), one closely related to *M. nudihortorum* (SAPN1) and one of unknown relationship (SAPN2).

#### TAXONOMY

***Radulomycetaceae*** Leal-Dutra, Dentinger, G.W. Griff., **fam. nov**. (Fig. 2D-F)

##### MYCOBANK

MB831047

##### ETYMOLOGY

From the type genus *Radulomyces*.

##### DIAGNOSIS

Basidiome resupinate, effused, mostly adnate, ceraceous, hymenophore smooth, tuberculate, odontioid, raduloid or poroid. Hyphal system monomitic, generative hyphae with clamps, hyaline, thin- to slightly thick-walled. Cystidia absent. Basidia terminal clavate or other form if pleural, usually with 4-sterigmata and a basal clamp. Basidiospores ellipsoid to globose, hyaline, mostly smooth, thin- to slightly thick-walled, acyanophilous, inamyloid and nondextrinoid.

##### TYPE GENUS

*Radulomyces* M.P. Christ.

##### NOTES

*Radulomyces, Aphanobasidium* and *Radulotubus* are placed in Radulomycetaceae fam. nov. Larsson (2007) suggested that *Lepidomyces* Jülich has affinities to *Aphanobasidium* and could possibly be placed in Pterulaceae. However, no sequence data for the genus are available. *Lepidomyces* is described as bearing pleurobasidia as in *Aphanobasidium*, but also leptocystidia as in *Coronicium* and *Merulicium*. Given its morphological similarities to the *Aphanobasidium* and the *Coronicium* superclade, we suggest to keep *Lepidomyces* as incertae sedis until molecular data are available to confirm its phylogenetic position.

***Phaeopterula*** (Henn.) Sacc. & D. Sacc., Syll. fung. **17**: 201 (1905) (Fig. 1M-O)

##### BASIONYM

*Pterula* subgen. *Phaeopterula* Henn., in Warburg, Monsunia 1: 9 (1899) [1900].

##### TYPE SPECIES

*Phaeopterula hirsuta* (Henn.) Sacc. & D. Sacc.

##### UPDATED DESCRIPTION

Basidiomes Pteruloid solitary or gregarious, scarcely branched to almost bushy, monopodial and slightly symmetric, branches from light brownish pink or greyish to pale brown and stipe dark reddish to rusty brown. Stipe surface glabrous with agglutinated hyphae (not sclerotioid) to villose-tomentose. Dark brown mycelial cords usually present. Hyphal system dimitic with thick-walled skeletal hyphae, generative hyphae thin-walled and often clamped. Hymenial cystidia absent, caulocystidia sometimes present. Basidia terminal, clavate to suburniform. Basidiospores less than 9 µm varying between pip-shaped, subamygdaliform and ellipsoid. Growing on dead twigs or dead wood.

##### NOTES

Hennings (1900) created the subgenus *Phaeopterula* to accommodate *Pterula hirsuta* that was distinguished from other *Pterula* spp. by its reportedly brown spores. Hennings (1904) later described a second species in the subgenus, *Ph. juruensis*, but noted that it was morphologically quite distinct from *Ph. hirsuta. Phaeopterula* was raised to genus level by Saccardo and Saccardo (1905) who cited only *Ph. juruensis. Pterula hirsuta* was recombined in *Dendrocladium* by Lloyd (1919) but later put back in *Pterula* by Corner (1950), even though Corner did not confirm the presence of brown spores in the samples he examined. Although we also have not observed pigmented spores in any of these taxa, dark brown pigments in the stipe hyphae are a consistent and diagnostic feature in this group, so we resurrect the name *Phaeopterula*. The term ‘Phaeo’ originally related to brown-pigmented basidiospores. Whilst members of this genus do not have brown basidiospores, they do contain brown hyphal pigments.

***Pterulicium*** Corner, Monograph of *Clavaria* and allied Genera: 689, 699 (1950) (Fig. 1I-L)

##### TYPE SPECIES

*Pterulicium xylogenum* (Berk. & Broome) Corner

##### UPDATED DESCRIPTION

Basidiomes pteruloid rarely corticioid, solitary or gregarious, simple or scarcely branched, occasionally exhibiting abundant unilateral branching (Fig. 1I and 1L), with colour varying from creamy white to brown on the stipe and creamy white on the tips or creamy white or pale lilaceous to pale brown on uniformly coloured basidiomes. Stipe surface sometimes sclerotioid [see Corner, (1950)]. Hyphal system dimitic with slightly thick-walled skeletal hyphae, generative hyphae thin-walled and often clamped. Hymenial cystidia usually present, caulocystidia sometimes present. Basidia terminal, clavate to suburniform. Basidiospores shape varying between globose to subglobose, pip-shaped, amygdaliform to subamygdaliform, ellipsoid. Growing on dead leaves, dead twigs or dead wood, rarely as a pathogen or endophyte of living plants.

##### NOTES

*Deflexula* is synonymised with *Pterulicium* in this study. In addition, several species previously placed in *Pterula* are transferred to *Pterulicium* (see the new combinations below). Other *Pterula* species that might need to be recombined in *Pterulicium*, require further investigation since their original descriptions do not provide enough information to confidently assign them here.

***Myrmecopterula*** Leal-Dutra, Dentinger & G.W. Griff., **gen. nov**. (Fig. 1A-F)

##### MYCOBANK

MB831048

##### ETYMOLOGY

From the ancient Greek word μύρμηκος (=mýrmēkos), genitive form of μύρμηξ (=mýrmēx), ants. Thus, *Pterula* of the ants, due to the observed relationship of several taxa in this genus with nests of fungus-growing ants.

##### TYPE SPECIES

*Myrmecopterula moniliformis* (Henn.) Leal-Dutra, Dentinger & G.W. Griff.

##### DIAGNOSIS

Usually associated with the nests of ants, growing on top or from living or dead nest or being cultivated by the ants. Bushy pteruloid basidiome, white-cream to light-brown and greyish surface, normally concolorous or stipe with a darker tone than the hymenophore, arising from cottony subiculum with mycelial cords, stipe surface sterile, dimitic hyphal system, relatively small spores (usually less than 7µm wide), or no basidiome. Differs from *Pterula* by the presence of the cottony subiculum.

##### NOTES

Basidiomes of *Myrmecopterula* species are very similar to *Pterula* in habit, shape and colour, but they differ in the presence of mycelial cords and of a cottony subiculum from which basidiomes emerge. Some species of *Myrmecopterula* arise from soil, while others superficially appear to grow on wood. Closer observation of wood-dwelling basidiomes revealed that rather than being lignicolous, instead they grow from a loose, granular substrate within a cavity inside the wood. This substrate in some cases resembles the substrate in the fungus gardens of *Apterostigma pilosum* group ants. In addition, *M. moniliformis*, which arises from soil, has been found emerging from active and inactive attine nests, (S. Sourell, pers. comm.; M.C. Aime, pers. comm.). Thus, all but one of the *Myrmecopterula* clades found to date had some association with attine ants, of which the two farmed mutualist species (*M. nudihortorum* and *M. velohortorum*) are best known. The five other species (of which only *M. moniliformis* is named) are less well studied and may play a role in decomposition of residual substrates in abandoned fungus garden, or potentially even as mycoparasites of the ant cultivar. In contrast, no *Pterula* spp. have any reported association with ants, but instead are found growing directly from wood and leaf litter.

***Myrmecopterula moniliformis*** (Henn.) Leal-Dutra, Dentinger & G.W. Griff., **comb. nov**. (Fig. 1E)

##### MYCOBANK

MB831049

##### BASIONYM

*Lachnocladium moniliforme* Henn., Hedwigia 43(3): 198 (1904).

##### SYNONYM

*Pterula moniliformis* (Henn.) Corner, Ann. Bot., Lond., n.s. 16: 569 (1952). *Thelephora clavarioides* Torrend, Brotéria, sér. bot. 12(1): 61 (1914).

Description in (Corner 1952b).

***Myrmecopterula nudihortorum*** (Dentinger) Leal-Dutra, Dentinger & G.W. Griff., **comb. nov** (Fig. 1D)

##### MYCOBANK

MB831050

##### BASIONYM

*Pterula nudihortorum* Dentinger [as ‘nudihortus’, later referred to as ‘nudihorta’], Index Fungorum 98: 1 (2014).

##### UPDATED DESCRIPTION

In the field, it is recognized by the absence of any veil on the fungus garden in the *Apterostigma* nests, usually inside decomposing trunks or underground. In culture, it forms very little aerial mycelium and exhibits very slow growth (2-3 mm/week radial growth rate on PDA at 25C). Hyphal clamps abundant.

##### NOTES

This species was formerly known as the ant cultivar G4. It is only known from the nest of fungus-growing ants in the *Apterostigma pilosum* group in the *A. manni* subclade (Schultz 2007).

***Myrmecopterula velohortorum*** (Dentinger) Leal-Dutra, Dentinger & G.W. Griff., **comb. nov**. (Fig. 1A)

##### MYCOBANK

MB831051

##### BASIONYM

*Pterula velohortorum* Dentinger [as ‘velohortus’, later referred to as ‘velohorta’], Index Fungorum 98: 1 (2014).

##### UPDATED DESCRIPTION

In the field, it is recognized by the *Apterostigma* garden covered by a mycelial veil, usually inside decomposing trunks, below the leaf litter or hanging on exposed surfaces aboveground. In culture, it forms very cottony aerial mycelia with presence of racquet hyphae (Fig. 5 in Additional file 3). Large and abundant hyphal clamps. Slow growth rate, but faster than *M. nudihortorum*.

##### NOTES

This species was formerly known as the ant cultivar G2. It is only known from the nest of fungus-growing ants in the *Apterostigma pilosum* group in the *A. dentigerum* subclade (Schultz 2007).

## DISCUSSION

### Introduction of Radulomycetaceae

We consider that it is better to erect a new family for these three genera (i.e. *Radulomyces, Radulotubus and Aphanobasidium*) than to leave them in Pterulaceae where they are clearly phylogenetically and morphologically distinct from nearly all the other members of Pterulaceae. In contrast, *Merulicium* (Fig. 2B-C) and *Coronicium* (Fig. 2A) form corticioid basidiomes but our phylogenetic analyses place them clearly within Pterulaceae. Two *Pterulicium* species, *Pm. echo* and *Pm. xylogenum*, also form both pteruloid and corticioid basidiomes, either independently or together (McLaughlin and McLaughlin 1980; Corner 1950).

Whilst the corticioid basidiomes of *Merulicium* and *Pm. echo* contain a dimitic hyphal system, typical of Pterulaceae, those of *Coronicium* spp. and *Pterulicium xylogenum* form a monomitic hyphal system, like all members of Radulomycetaceae. However, no members of Radulomycetaceae form cystidia, whereas these cells are found in most Pterulaceae (McLaughlin and McLaughlin 1980; Corner 1970, 1967, 1952a, b, 1950; Bernicchia and Gorjón 2010), including *Coronicium* spp. Thus, Radulomycetaceae fam. nov. is morphologically characterized by the combination of resupinate basidiomes, monomitic hyphal system and lack of cystidia. Moreover, our phylogenetic analyses strongly support the segregation of Radulomycetaceae fam. nov. from Pterulaceae.

### Reintroduction of Phaeopterula

*Phaeopterula* spp. are distinct from other pterulaceous genera due to the distinctive brown colour of the main axis of the basidiome and monopodial/symmetric branching of these structures. This contrasts with other Pterulaceae which are either highly branched (bushy) and of uniform colour (*Pterula* and *Myrmecopterula*) or pigmented only at the stipe base, and (mostly) unbranched (*Pterulicium*). Hennings (1900) originally defined *Phaeopterula* by its brown spores. Corner (1950) cast doubt on the significance of this trait, but our results show that, despite an apparently misguided justification, Hennings was correct to group *Ph. juruensis* with *Ph. hirsuta*.

All *Phaeopterula* spp. are exclusively found on decaying wood, whereas members of other genera of Pterulaceae inhabit more diverse lignocellulosic substrates. Given the basal position of *Phaeopterula* in Pterulaceae, and the fact that all members of the sister family Radulomycetaceae are also lignicolous on wood, this habit is parsimoniously the ancestral condition. The reintroduction of *Phaeopterula* aims to pay tribute to Paul Hennings’ work and his contribution to the taxonomy of Pterulaceae.

### Synonymy of Deflexula with Pterulicium

Besides the paraphyly represented by *Phaeopterula*, the *Pterulicium* clade shows the clear polyphyly of *Pterula* and *Deflexula*. Several species in the two latter genera are intermixed in a strongly supported subclade (Fig. 3). The presence of the type species of both *Deflexula* and *Pterulicium* within this clade requires that only one name be kept. Both genera were proposed by Corner (1950), to accommodate the dimitic and coralloid (but non-bushy) species, not fitting the description of *Pterula*. The name *Pterulicium* was based on a ‘portmanteau’ combination of *Pterula* and *Corticium* to reflect the presence of a corticioid patch at the stipe base (Corner 1950). However, this patch has only been reported in two species, *Pterulicium xylogenum* (Corner 1950) and *Pm. echo* (McLaughlin and McLaughlin 1980). *Deflexula* was named for the downward-oriented (positively geotropic) basidiomes (Corner 1950). Corner (1950) stated that the resupinate patch in *Pterulicium xylogenum* is monomitic, can exist independently of the coralloid basidiome and is fertile when facing downward; he suggested that there was a close similarity between *Deflexula* and *Pterulicium* in the way the resupinate patch develops from the base of the basidiome. He also made a case for the formation of a fertile hymenium when facing downward in the two genera as supporting this similarity. Nonetheless, experimental studies on *Pm. echo* show that orientation of the hymenium does not affect the ability to produce spores, i.e., the hymenium is ageotropic (McLaughlin et al. 1978) and raised doubts about the validity of the genus *Deflexula*. This morphological distinction is not supported by phylogenetic analysis (Dentinger et al. 2009, Fig. 3) and its emphasis through taxonomic preservation would perpetuate misunderstanding. Accordingly, we propose to retain *Pterulicium* for this clade to avoid major misinterpretations of the species morphology.

### Introduction of Myrmecopterula gen. nov

Two species of Pterulaceae are cultivated by fungus-farming ants of the *Apterostigma pilosum* group in South and Central America (Dentinger et al. 2009; Munkacsi et al. 2004; Villesen et al. 2004; Mueller et al. 2018). Despite intensive investigation, neither has been observed to form basidiomes, but *M. velohortorum* is characterised by the formation of a veil of mycelium around the fungus garden, whilst *M. nudihortorum* lacks this veil. We recovered both species in a strongly supported clade, as a sister clade of *Pterula*, alongside five other subclades containing fertile, apparently free-living species.

All the samples in this clade were collected from neotropical habitats (Fig. 1A-F), mostly as part of our recent fieldwork. During sampling campaigns by ourselves and others, it was observed that many of the ‘free-living’ specimens were associated in some way with living ant colonies or abandoned attine nests. Two *Myrmecopterula* samples belonging to subclade SAPV1 (CALD170307-02 and CALD170307-03; Fig. 1A) were found forming basidiomes atop two distinct but adjacent (1 m apart) living *Apterostigma* nests in Amazonian Rainforest. The cultivated mutualists from both nests were also analysed and found to belong to *M. velohortorum* confirming that the basidiomes were not linked to the cultivated mycelia in these nests. The third member of subclade SAPV1 was also reported forming a nascent basidiome on a living Apterostigma nest in Panama (Munkacsi et al. 2004). *M. moniliformis* (SAPN1; Fig. 1E) has been reported to be found outside both active and apparently inactive (see *Myrmecopterula*: Notes on Taxonomy section above) attine nests (S. Sourell, pers. comm.; M.C. Aime, pers.comm.) as was CALD170315-04 (SAPV2; Fig. 1B) and CALD170122-04 (SAPV3; Fig. 1C). Lastly, the mycelium of one sample (JSP 07-03 B 5.1; SAPV3) was isolated from a living *Atta capiguara* nest by Pereira et al. (2016).

The observations above and the phylogenetic analyses suggests that association with attine ants is a widespread trait amongst members of this clade, hence its naming as *Myrmecopterula*.

Most recent attention on Pterulaceae has been lavished on the ant-cultivated mutualists *M. nudihortorum* and *M. velohortorum*. These were once thought to be sister clades (Munkacsi et al. 2004; Villesen et al. 2004) but are now known to be only distantly related within the *Myrmecopterula* clade (Dentinger et al. 2009, Fig. 3). This suggests two possibilities for the evolution of the *Myrmecopterula-Apterostigma* mutualism: 1) that it evolved independently on two occasions or, 2) that it is an ancestral condition of all *Myrmecopterula*. However, it is at present unclear whether the extant mutualistic association found for *M. nudihortorum* and *M. velohortorum* is ancestral, implying that the other taxa escaped the mutualism, or whether the looser association with ant nests widespread amongst members of *Myrmecopterula* was more recently elevated to a higher level of interdependence for these two species, as suggested by Dentinger et al. (2009). It is also possible that the free-living species within the *Myrmecopterula* may be specialised parasites specifically targeting their sister species that have formed a mutualism with the ants. An analogous situation is found in the leaf-cutting ants species *Acromyrmex echinatior* and its sister species *Acromyrmex insinuator*, the latter a highly specialised social parasite of the former (Sumner et al. 2004).

The basis of the association of ‘free-living’ species with attine ants and/or their abandoned nests is unclear. Given the apparent preference of some for abandoned nests, they may be specialised early stage colonisers of ant nest debris. A further possibility is that they are cheaters, deriving nutrition from the ant-collected biomass but not reciprocating by producing hyphae palatable to ants. This would represent a novel form of fungal mimicry, perhaps achieved by the ants’ inability to differentiate hyphae of closely related species. Lastly, they may be mycoparasitic, including on ant cultivars, although there is currently no direct evidence supporting this hypothesis.

### Re-delimitation of Pterulaceae

All the accepted genera in Pterulaceae were sampled in this study except for the monotypic *Allantula*. One specimen, with morphology consistent with Corner’s description of *Allantula diffusa*, with pteruloid basidiomes borne on slender mycelial cords as curved intercalary swellings, was collected during our fieldwork (Fig. 1M). Phylogenetic reconstruction placed this specimen firmly within *Phaeopterula*. However, we have been unable to obtain the type specimen (no other collections authenticated exist) for more detailed analysis.

Thus, we re-delimit Pterulaceae containing six genera: *Allantula, Coronicium, Merulicium, Myrmecopterula, Phaeopterula, Pterula and Pterulicium*.

## CONCLUSION

In this study, we presented a reclassification of Pterulaceae based on morphological and phylogenetic analyses with samples from six out of seven genera previously accepted in the family. Three early diverging resupinate genera were placed in Radulomycetaceae fam. nov. (*Aphanobasidium, Radulomyces and Radulotubus*), *Myrmecopterula* gen. nov. was erected to accommodate ant associated species previously classified in *Pterula*; several species from the latter were also recombined in the reintroduced *Phaeopterula* and in *Pterulicium*, and finally *Deflexula* was synonymised with *Pterulicium*. Pterulaceae was thus redelimited to accommodate seven genera *Allantula, Coronicium, Merulicium, Myrmecopterula, Phaeopterula, Pterula* and *Pterulicium*. Some species kept in *Pterula* might also need to be recombined since the original description was not enough to make these changes. Type specimens should be analysed considering the delimitations proposed in this study.

## Supporting information

Suppdata1

Suppdata2

Suppdata3

Suppdata4

## DECLARATIONS

### Ethics approval and consent to participate

Not applicable

### Adherence to national and international regulations

All the requirements for specimen acquisition, transportation and study in Brazil, were followed according to the Brazilian federal regulations. Dried samples were transferred between fungaria following the regulations of the Nagoya Protocol to the Convention on Biological Diversity 2011.

### Consent for publication

Not applicable

### Availability of data and material

Details of the availability of the data and material used in this study can be found within the text.

DNA sequences were submitted to NCBI Genbank database (see Table 2 and Additional file 1). Alignments were deposited at Tree-Base (ID: 24428). Dried specimens are deposited in the fungaria listed the Methods section.

### Authors’ contributions

MAN, BTMD and CALD conceived the study; CALD and MAN obtained permits for fieldwork in Brazil; all authors carried sample collection; CALD, LAC and BTMD performed molecular methods; CALD performed phylogenetic analyses; CALD, GWG and BTMD drafted the manuscript; all authors approved the final version of the manuscript.

## Acknowledgements

This article is part of the PhD thesis of CALD. The authors are grateful to the managers of Parque Nacional da Tijuca (ICMBio), Floresta Nacional do Tapajós, Parque Nacional do Iguaçú, Parque Nacional da Serra dos Orgãos and Reserva Biológica Augusto Ruschi, for logistical support and collection permits. Biological Dynamics of Forest Fragmentation Project (BDFFP) for providing logistical and field support. This is publication XXX of the BDFFP - INPA/STRI Technical Series (field reserve; location of specimen used here). We acknowledge the support of the Supercomputing Wales project, which is part-funded by the European Regional Development Fund (ERDF) via Welsh Government. We are also thankful for the kind support and loan from the following herbaria: FLOR (Universidade Federal de Santa Catarina), K (Royal Botanical Gardens, Kew), INPA (Instituto Nacional de Pesquisas da Amazônia), HSTM (Universidade Federal do Oeste do Pará). The researchers who kindly collect and provide samples or photos, or helped in other way to this study: Ted Schultz, M. Catherine Aime, Dennis Desjardin, D. Jean Lodge, Bitty Roy, Michael Wherley, Tobias Policha, Jesse McAlpine, Tommy Jenkinson, Rocío Manobanda, Celeste Heisecke, Altielys C. Magnago, Eduardo P. Fazolino, Ariadne N. M. Furtado, Emerson L. Gumboski, Elisandro R. Drechsler-Santos, Julia Simon, Cauê Oliveira, Stefan Blaser, David J. Harries, Genivaldo Alves da Silva, Susanne Sourell, Lucie Zíbarová, André Rodrigues, Quimi Vidaurre Montoya, Louro Lima, Ocírio Juruna de Souza Pereira, Laura Martínez-Suz, Shaun Pennycook, Phillip Camarota Moura and Rafael Trevisan.

## Key to genera of Pterulaceae and Radulomycetaceae fam. nov

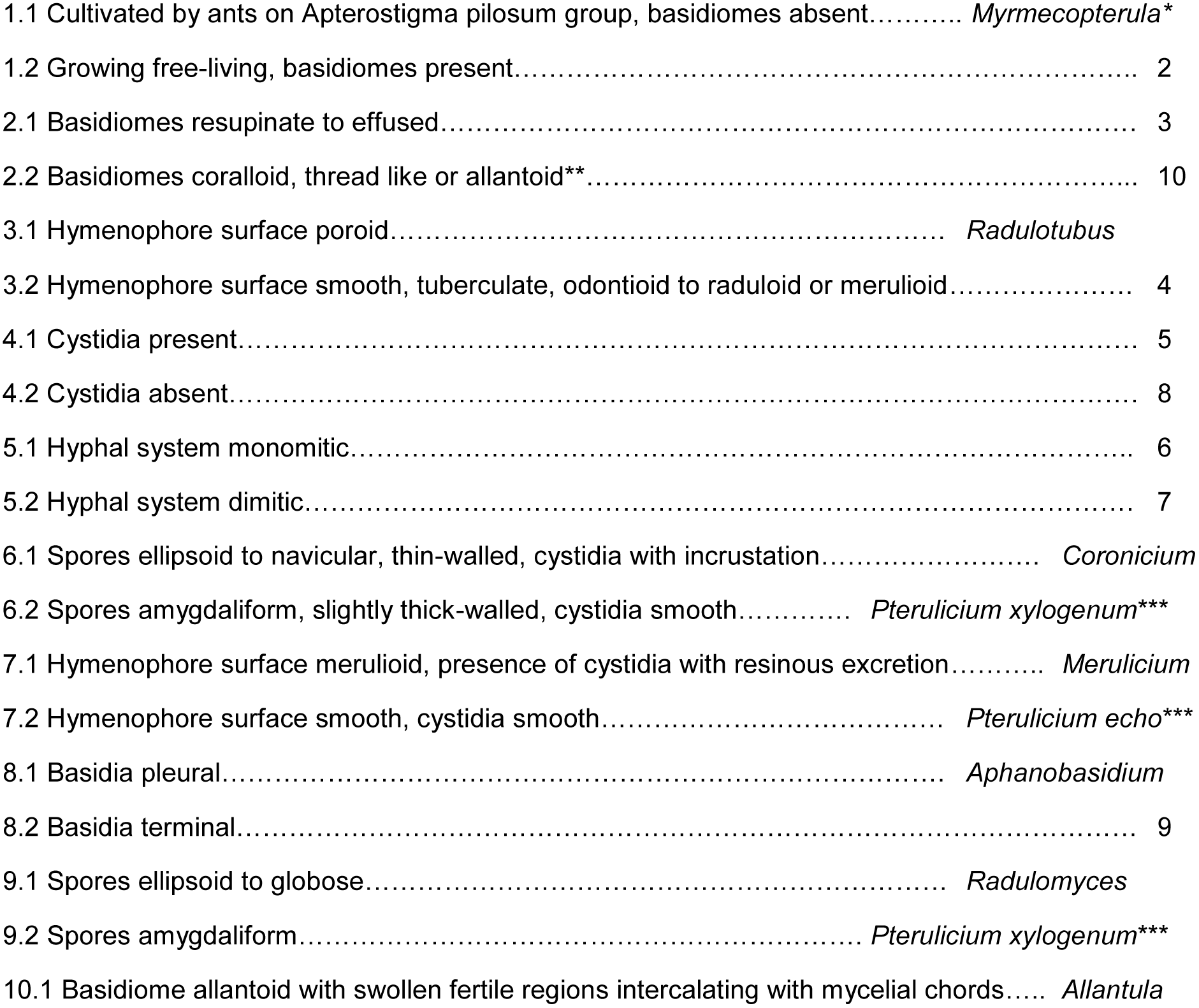
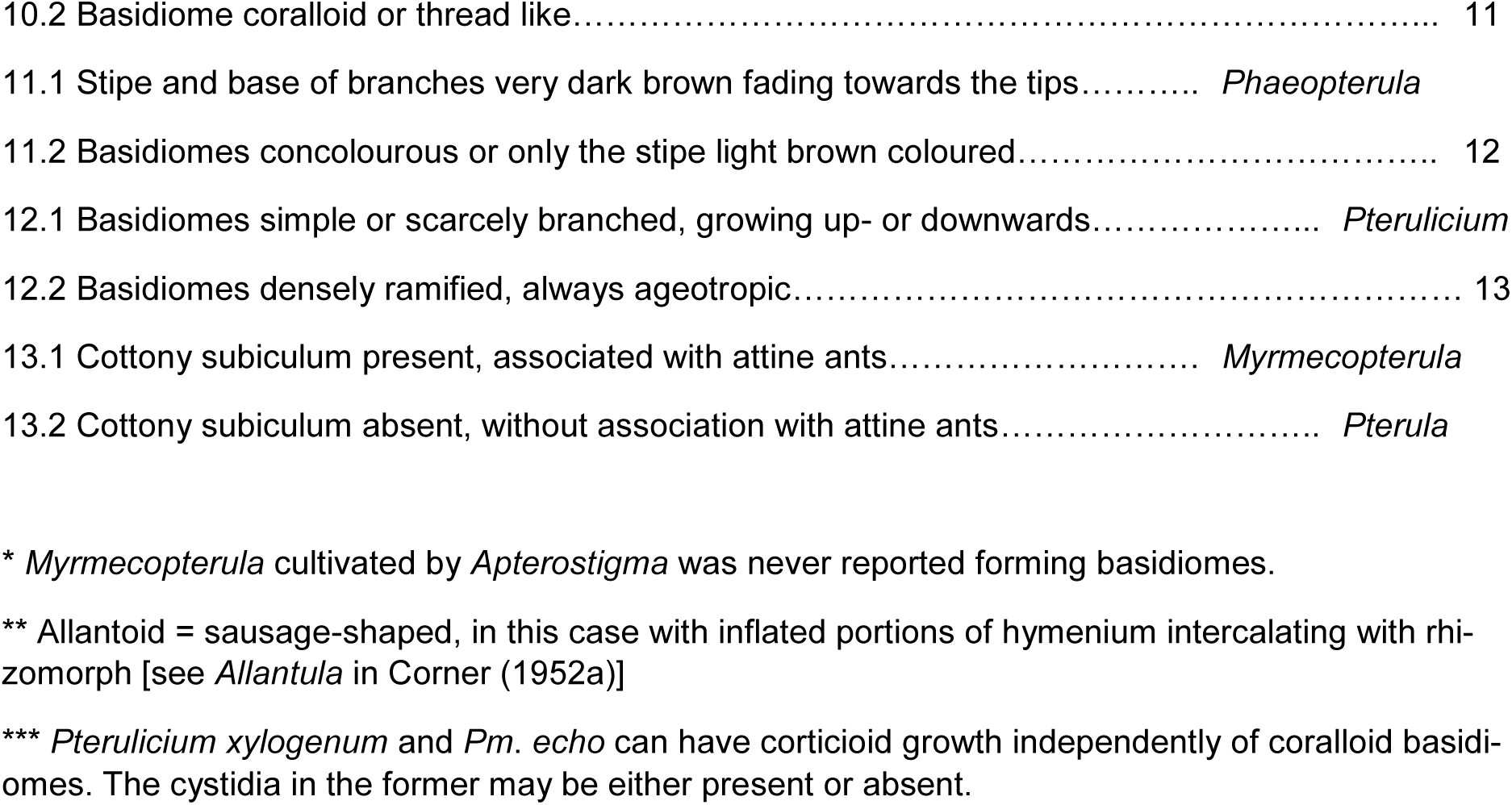

## ADDITIONAL NOMENCLATURAL NOVELTIES

***Phaeopterula anomala*** (P. Roberts) Leal-Dutra,

Dentinger, G.W. Griff., **comb. nov**.

MycoBank MB830999

Basionym: *Pterula anomala* P. Roberts, *Kew*

*Bull*. **54**(3): 528 (1999).

Description in Roberts (1999).

***Phaeopterula hirsuta*** (Henn.) Sacc. & D. Sacc.,

*Syll. fung*. (Abellini) **17**: 201 (1905)

MycoBank MB469044

Basionym: *Pterula hirsuta* Henn., in Warburg,

*Monsunia* **1**: 9 (1899) [1900].

Synonym: *Dendrocladium hirsutum* (Henn.)

Lloyd, *Mycol. Writ*. **5**: 870 (1919).

Description in Corner (1950).

***Phaeopterula juruensis*** Henn. ex Sacc. & D. Sacc., *Syll. fung*. (Abellini) **17**: 201 (1905) **(Fig. 1O)**

MycoBank MB634235

Basionym: *Pterula juruensis* Henn. [as ‘*Phaeopterula juruensis*’], *Hedwigia* **43**(3): 175 (1904).

Synonym: *Dendrocladium juruense* (Henn.)

Lloyd, *Mycol. Writ*. **5**: 870 (1919).

Descriptions in Corner (1950, 1952b).

***Phaeopterula stipata*** (Corner) Leal-Dutra, Dentinger, G.W. Griff., **comb. nov. (Fig. 1N)**

MycoBank MB831000

Basionym: *Pterula stipata* Corner, *Ann. Bot*., Lond., n.s. **16**: 568 (1952).

Description in Corner (1952b).

***Phaeopterula taxiformis*** (Mont.) Leal-Dutra, Dentinger, G.W. Griff., **comb. nov**.

MycoBank MB831001

Basionym: *Pterula taxiformis* Mont., *Syll. gen. sp. crypt*. (Paris): 181 (1856).

Synonyms: *Lachnocladium taxiforme* (Mont.) Sacc., *Syll. fung*. (Abellini) **6**: 740 (1888). *Pterula humilis* Speg., *Revista Argent. Hist. Nat*. **1**(2): 110 (1891). *Pterula humilis* var. *tucumanensis* Speg., *Anal. Mus. nac. B. Aires*, Ser. 3 **12**: 280 (1909).

Descriptions in Corner (1950, 1952b).

***Phaeopterula taxiformis* var. *gracilis*** (Corner) Leal-Dutra, Dentinger, G.W. Griff., **comb. nov**.

MycoBank MB831002

Basionym: *Pterula taxiformis* var. *gracilis* Corner,

*Ann*. *Bot*., Lond., n.s. 16: 568 (1952).

Description in Corner (1952b).

***Pterulicium argentinum*** (Speg.) Leal-Dutra, Dentinger, G.W. Griff., **comb. nov**.

MycoBank MB831003

Basionym: *Mucronella argentina* Speg., *Anal. Mus. nac. Hist. nat. B. Aires* **6**: 178 (1898) [1899].

Synonyms: *Deflexula argentina* (Speg.) Corner, *Ann. Bot*., Lond., n.s. **16**: 276 (1952). *Deflexula lilaceobrunnea* var. *elongata* Corner, *Ann. Bot*.,

Lond., n.s. **16**: 276 (1952).

Description in Corner (1952a, 1970)

***Pterulicium argentinum* var. *ramosum*** (Corner) Leal-Dutra, Dentinger, G.W. Griff., **comb. nov**.

MycoBank MB831004

Basionym: *Deflexula argentina* (Speg.) Corner,

*Ann. Bot*., Lond., n.s. **16**: 276 (1952).

Description in Corner (1970).

***Pterulicium bambusae*** (Corner) Leal-Dutra, Dentinger, G.W. Griff., **comb. nov**.

MycoBank MB831005

Basionym: *Pterula bambusae* Corner, *Beih Nova*.

*Hedwigia* **33**: 209 (1970).

Description in Corner (1970).

***Pterulicium bromeliphilum*** (Corner) Leal-Dutra,

Dentinger, G.W. Griff., **comb. nov**.

MycoBank MB831006

Basionym: *Pterula bromeliphil*a Corner, *Beih*.

*Nova Hedwigia* **33**: 210 (1970).

Description in Corner (1970).

***Pterulicium brunneosetosum*** (Corner) Leal-Dutra, Dentinger, G.W. Griff., **comb. nov**.

MycoBank MB831007

Basionym: *Pterula brunneosetosa* Corner, *Ann*.

*Bot*., Lond., n.s. **16**: 566 (1952)

Descriptions in Corner (1952b, 1957, 1970).

***Pterulicium campoi*** (Speg.) Leal-Dutra, Dentinger, G.W. Griff., **comb. nov**.

MycoBank MB831008

Basionym: *Pterula campoi* Speg., *Bol. Acad. nac*.

*Cienc. Córdoba* **25**: 29 (1921).

Descriptions in Corner (1970) and Spegazzini (1921).

***Pterulicium caricis-pendulae*** (Corner) Leal-Dutra, Dentinger, G.W. Griff., **comb. nov**.

MycoBank MB831009

Basionym: *Pterula caricis-pendulae* Corner, *Beih*.

*Nova Hedwigia* **33**: 211 (1970).

Description in Corner (1970).

***Pterulicium crassisporum*** (P. Roberts) Leal-Dutra, Dentinger, G.W. Griff., **comb. nov**.

MycoBank MB831010

Basionym: *Pterula crassispora* P. Roberts, *Kew*

*Bull*. **54**(3): 531 (1999).

Description in Roberts (1999).

***Pterulicium cystidiatum*** (Corner) Leal-Dutra, Dentinger, G.W. Griff., **comb. nov**.

MycoBank MB831011

Basionym: *Pterula cystidiata* Corner, *Ann. Bot*.,

Lond., n.s. **16**: 567 (1952).

Description in Corner (1952b).

***Pterulicium debile*** (Corner) Leal-Dutra, Dent-inger, G.W. Griff., **comb. nov**.

MycoBank MB831012

Basionym: *Pterula bromeliphil*a Corner, *Monograph of Clavaria and allied Genera*, (Annals of Botany Memoirs No. 1): 698 (1950).

Description in (Corner 1950).

***Pterulicium echo*** (D.J. McLaughlin & E.G. McLaughlin) Leal-Dutra, Dentinger, G.W. Griff., **comb. nov**.

MycoBank MB831013

Basionym: *Pterula echo* D.J. McLaughlin & E.G.

McLaughlin, *Can. J. Bot*. **58**: 1328 (1980).

Description in McLaughlin and McLaughlin (1980).

***Pterulicium epiphylloides*** (Corner) Leal-Dutra, Dentinger, G.W. Griff., **comb. nov**.

MycoBank MB831014

Basionym: *Pterula epiphylloides* Corner, *Ann*.

*Bot*., Lond., n.s. **16**: 567 (1952).

Description in Corner (1952b).

***Pterulicium epiphyllum*** (Corner) Leal-Dutra, Dentinger, G.W. Griff., **comb. nov**.

MycoBank MB831015

Basionym: *Pterula epiphylla* Corner, *Monograph of Clavaria and allied Genera*, (Annals of Botany Memoirs No. 1): 698 (1950).

Description in (Corner 1950).

***Pterulicium fasciculare*** (Bres. & Pat.) Leal-Dutra, Dentinger, G.W. Griff., **comb. nov**.

MycoBank MB831016

Basionym: *Pterula fascicularis* Bres. & Pat., *Mycol. Writ*. **1**: 50 (1901).

Synonym: *Deflexula fascicularis* (Bres. & Pat.) Corner, *Monograph of Clavaria and allied Genera*, (Annals of Botany Memoirs No. 1): 395 (1950).

Description in Corner (1950).

***Pterulicium fluminense*** (Corner) Leal-Dutra, Dentinger, G.W. Griff., **comb. nov**.

MycoBank MB831017

Basionym: *Pterula fluminensis* Corner, *Ann. Bot*.,

Lond., n.s. **16**: 567 (1952).

Descriptions in Corner (1952b, 1970).

***Pterulicium gordium*** (Speg.) Leal-Dutra, Dentinger, G.W. Griff., **comb. nov**.

MycoBank MB831018

Basionym: *Clavaria gordius* Speg., *Anal. Soc*.

*cient. argent*. **17**(2): 83 (1884).

Synonym: *Pterula gordius* (Speg.) Corner, *Monograph of Clavaria and allied Genera*, (Annals of Botany Memoirs No. 1): 513 (1950).

Description in (Corner 1950).

***Pterulicium gordium* var. *macrosporum*** (Corner) Leal-Dutra, Dentinger, G.W. Griff., **comb. nov**.

MycoBank MB831019

Basionym: *Pterula gordius* var. *macrospora* Corner, *Proc. Linn. Soc*. London **178**: 100 (1967).

Description in Corner (1967).

***Pterulicium gracile*** (Desm. & Berk.) Leal-Dutra, Dentinger, G.W. Griff., **comb. nov**.

MycoBank MB831020

Basionym: *Typhula gracilis* Desm. & Berk., Ann. nat. Hist., Mag. Zool. Bot. Geol. 1: 202 (1838).

Synonyms: *Pistillaria gracilis* (Desm. & Berk.) Pat., *Tab. analyt. Fung*. (Paris)(6): 30 (1886). *Hirsutella gracilis* (Desm. & Berk.) Pat., *Revue mycol*., Toulouse **14**(no. 54): 69 (1892). *Pterula gracilis* (Desm. & Berk.) Corner, *Monograph of Clavaria and allied Genera*, (Annals of Botany Memoirs No. 1): 514 (1950). *Clavaria aculina* Quél., *C. r. Assoc. Franç. Avancem. Sci*. **9**: 670 (1881) [1880]. *Pistillaria aculina* (Quél.) Pat., *Tab. analyt. Fung*. (Paris)(6): 29 (fig. 570) (1886). *Ceratella aculina* (Quél.) Pat., *Hyménomyc. Eur*. (Paris): 157 (1887). *Cnazonaria aculina* (Quél.) Donk, *Meded. Bot. Mus. Herb. Rijks Univ*. Utrecht **9**: 97 (1933). *Pistillaria aculina* subsp. *juncicola* Bourdot & Galzin, *Hyménomyc. de France* (Sceaux): 138 (1928) [1927]. *Pistillaria aculina* subsp. *graminicola* Bourdot & Galzin, *Hyménomyc. de France* (Sceaux): 139 (1928) [1927]. *Pistillaria aculina* subsp. *acicula* Bourdot & Galzin, *Hyménomyc. de France* (Sceaux): 139 (1928) [1927]. *Typhula brunaudii* Quél., *C. r. Assoc. Franç. Avancem. Sci*. **13**: 283 (1885) [1884]. *Clavaria brunaudii* (Quél.) Sacc., *Syll. fung*. (Abellini) **6**: 730 (1888). *Ceratella ferryi* Quél. & Fautrey, *Revue mycol*., Toulouse **15**(no. 57): 15 (1893). *Pistillaria ferryi* (Quél. & Fautrey) Sacc., *Syll. fung*. (Abellini) **11**: 141 (1895). *Pistillaria ferryi* subsp. *tremula* Sacc., *Syll. fung*. (Abellini) **17**: 202 (1905). *Mucronella rickii* Oudem., *Ned*. *kruidk. Archf*, 3 sér. **2**(3): 667 (1902). *Cnazonaria rickii* (Oudem.) Donk, *Meded. Bot. Mus. Herb. Rijks Univ*. Utrecht **9**: 99 (1933). *Ceratellopsis rickii* (Oudem.) Corner, *Monograph of Clavaria and allied Genera*, (Annals of Botany Memoirs No. 1): 205 (1950).

Description in (Corner 1950).

***Pterulicium incarnatum*** (Pat.) Leal-Dutra, Dentinger, G.W. Griff., **comb. nov**.

MycoBank MB831021

Basionym: *Pterula incarnata* Pat., in Patouillard & Lagerheim, *Bull. Herb. Boissier* **3**(1): 58 (1895)

Description in (Corner 1950, 1970)

***Pterulicium intermedium*** (Dogma) Leal-Dutra, Dentinger, G.W. Griff., **comb. nov**.

MycoBank MB831022

Basionym: *Pterula intermedia* Dogma, *Philipp. Agric*. **49**: 852 (1966)

Descriptions in (Corner 1970) and Dogma (1966)

***Pterulicium laxum*** (Pat.) Leal-Dutra, Dentinger, G.W. Griff., **comb. nov**.

MycoBank MB831023

Basionym: *Pterula laxa* Pat., *Bull. Soc. mycol. Fr*. **18**(2): 175 (1902).

Description in (Corner 1950, 1970) Pat. 1902

***Pterulicium lilaceobrunneum*** (Corner) Leal-Dutra, Dentinger, G.W. Griff., **comb. nov. (Fig. 1K)**

MycoBank MB831024

Basionym: *Deflexula lilaceobrunnea* Corner, *Monograph of Clavaria and allied Genera*, (Annals of Botany Memoirs No. 1): 695 (1950).

Description in Corner (1952a).

***Pterulicium lilaceobrunneum* var. *evolutius*** (Corner) Leal-Dutra, Dentinger, G.W. Griff., **comb. nov**.

MycoBank MB831025

Basionym: *Deflexula lilaceobrunnea* var. *evolutior* Corner, *Beih. Nova*

*Hedwigia* **33**: 197 (1970).

Description in Corner (1970).

***Pterulicium longisporum*** (Corner) Leal-Dutra, Dentinger, G.W. Griff., **comb. nov**.

MycoBank MB831026

Basionym: *Pterula longispora* Corner, *Ann. Bot*.,

Lond., n.s. **16**: 567 (1952)

Description in (Corner 1952b).

***Pterulicium macrosporum*** (Pat.) Leal-Dutra, Dentinger, G.W. Griff., **comb. nov**.

MycoBank MB831027

Basionym: *Ceratella macrospora* Pat., in Patouillard & Lagerheim, *Bull. Soc. mycol. Fr*. **8**(3): 119 (1892).

Synonyms: *Pistillaria macrospora* (Pat.) Sacc., *Syll. fung*. (Abellini) **11**: 142 (1895). *Pterula macrospora* (Pat.) Corner, *Monograph of Clavaria and allied Genera*, (Annals of Botany Memoirs No. 1): 518 (1950).

Description in (Corner 1950, 1970). Pat 1892

***Pterulicium majus*** (Corner) Leal-Dutra, Dentinger, G.W. Griff., **comb. nov**.

MycoBank MB831028

Basionym: *Deflexula major* Corner, *Ann. Bot*., Lond., n.s. **16**: 277 (1952).

Description in Corner (1952a).

***Pterulicium mangiforme*** (Corner) Leal-Dutra, Dentinger, G.W. Griff., **comb. nov**.

MycoBank MB831029

Basionym: *Deflexula mangiformis* Corner, *Ann. Bot*., Lond., n.s. **16**: 278 (1952).

Description in Corner (1952a).

***Pterulicium microsporum*** (Corner) Leal-Dutra, Dentinger, G.W. Griff., **comb. nov**.

MycoBank MB831030

Basionym: *Deflexula microspora* Corner, *Bull*.

*Jard. bot. État Brux*. **36**: 264 (1966).

Description in Corner (1966).

***Pterulicium nanum*** (Pat.) Leal-Dutra, Dentinger, G.W. Griff., **comb. nov**.

MycoBank MB831031

Basionym: *Pterula nana* Pat., Bull. *Soc. mycol. Fr*. **18**(2): 175 (1902).

Synonyms: *Deflexula nana* (Pat.) Corner, *Bull. Jard. bot. État Brux*. **36**: 264 (1966). *Pterula vanderystii* Henn. [as *‘vanderysti’*], *Ann. Mus. Congo Belge*, Bot., Sér. 5 **2**(2): 96 (1907). *Deflexula vanderystii* (Henn.) Corner, *Ann. Bot*., Lond., n.s. **16**: 284 (1952).

Description in Corner (1966).

***Pterulicium naviculum*** (Corner) Leal-Dutra, Dentinger, G.W. Griff., **comb. nov**.

MycoBank MB831032

Basionym: *Pterula navicula* Corner, *Ann. Bot*., Lond., n.s. **16**: 568 (1952).

Description in (Corner 1952b).

***Pterulicium oryzae*** (Remsberg) Leal-Dutra, Dentinger, G.W. Griff., **comb. nov**.

MycoBank MB831033

Basionym: *Pistillaria oryzae* Remsberg, *Mycologia* **32**(5): 668 (1940).

Synonym: *Pterula oryzae* (Remsberg) Corner, *Monograph of Clavaria and allied Genera*, (Annals of Botany Memoirs No. 1): 519 (1950).

Descriptions in (Corner 1950) and Remsberg (1940)

***Pterulicium phyllodicola*** (Corner) Leal-Dutra, Dentinger, G.W. Griff., **comb. nov**.

MycoBank MB831034

Basionym: *Pterula phyllodicola* Corner, *Beih. Nova Hedwigia* **33**: 220 (1970).

Description in Corner (1970)

***Pterulicium phyllophilum*** (McAlpine) Leal-Dutra, Dentinger, G.W. Griff., **comb. nov**.

MycoBank MB831035

Basionym: *Clavaria phyllophila* McAlpine, *Agric. Gaz. N*.*S*.*W*., Sydney **7**: 86 (1896).

Synonym: *Pterula phyllophila* (McAlpine) Corner, *Monograph of Clavaria and allied Genera*, (Annals of Botany Memoirs No. 1): 520 (1950).

Description in (Corner 1950).

***Pterulicium rigidum*** (Donk) Leal-Dutra, Dentinger, G.W. Griff., **comb. nov**.

MycoBank MB831036

Basionym: *Pterula rigida* Donk, *Monograph of Clavaria and allied Genera*, (Annals of Botany Memoirs No. 1): 698 (1950).

Description in Corner (1950)

***Pterulicium sclerotiicola*** (Berthier) Leal-Dutra, Dentinger, G.W. Griff., **comb. nov**.

MycoBank MB831037

Basionym: *Pterula sclerotiicola* Berthier, *Bull*.

*trimest. Soc. mycol. Fr*. **83**: 731 (1968) [1967]

Description in Corner (1970)

***Pterulicium secundirameum*** (Lév) Leal-Dutra, Dentinger, G.W. Griff., **comb. nov. (Fig. 1I)**

MycoBank MB831038

Basionym: *Clavaria secundiramea* Lév., *Annls Sci. Nat*., *Bot*., sér. 3 **2**: 216 (1844).

Synonyms: *Pterula secundiramea* (Lév.) Speg., *Bol. Acad. nac. Cienc. Córdoba* **11**(4): 466 (1889). *Deflexula secundiramea* (Lév.) Corner, *Beih. Nova Hedwigia* **33**: 199 (1970). *Pterula palmicola* Corner, *Ann. Bot*., Lond., n.s. **16**: 568 (1952).

Descriptions in Corner (1950, 1952b).

Notes

The synonymisation of *Pm. palmicola* (samples M50 and M83) in *Pm. secundirameum* (samples M70 and genome5) is based on our phylogenetic results and morphological comparisons. The only morphological difference between the two species is the shape of the basidiome, however, the other characters are similar and both species are nested together within our tree (Additional file 2).

***Pterulicium sprucei*** (Mont.) Leal-Dutra, Dentinger, G.W. Griff., **comb. nov. (Fig. 1L)**

MycoBank MB831039

Basionym: *Hydnum sprucei* Mont., *Syll. gen. sp. crypt*. (Paris): 173 (1856).

Synonyms: *Pterula sprucei* (Mont.) Lloyd, *Mycol. Writ*. **5**: 865 (1919). *Deflexula sprucei* (Mont.) Maas Geest., *Persoonia* **3**(2): 179 (1964). *Pterula pennata* Henn., *Hedwigia* **43**(3): 174 (1904). *Deflexula pennata* (Henn.) Corner, *Ann. Bot*., Lond., n.s. **16**: 278 (1952).

Description in Corner (1952a) as *D. pennata*, Corner (1970) and Maas Geesteranus (1964).

***Pterulicium subsimplex*** (Henn.) Leal-Dutra, Dentinger, G.W. Griff., **comb. nov**.

MycoBank MB831040

Basionym: *Pterula subsimplex* Henn., *Hedwigia* **36**(4): 197 (1897).

Synonyms: *Deflexula subsimplex* (Henn.) Corner, *Ann. Bot*., Lond., n.s. **16**: 279 (1952). *Pterula nivea* Pat., *Bull. Soc. mycol. Fr*. **18**(2): 174 (1902). *Deflexula nivea* (Pat.) Corner, *Monograph of Clavaria and allied Genera*, (Annals of Botany Memoirs No. 1): 398 (1950). *Mucronella pacifica* Kobayasi, *Bot. Mag*., Tokyo **53**: 160 (1939). *Deflexula pacifica* (Kobayasi) Corner, *Monograph of Clavaria and allied Genera*, (Annals of Botany Memoirs No. 1): 399 (1950).

Descriptions in Corner (1952a) and Corner (1950) as D. pacifica.

***Pterulicium subsimplex* var. *multifidum*** (Corner) Leal-Dutra, Dentinger, G.W. Griff., **comb. nov**.

MycoBank MB831041

Basionym: *Deflexula subsimplex* var. *multifida*

Corner, *Ann. Bot*., Lond., n.s. **16**: 282 (1952).

Description in Corner (1952a).

***Pterulicium subtyphuloides*** (Corner) Leal-Dutra, Dentinger, G.W. Griff., **comb. nov**.

MycoBank MB831042

Basionym: *Pterula subtyphuloides* Corner, *Monograph of Clavaria and allied Genera*, (Annals of Botany Memoirs No. 1): 698 (1950).

Description in Corner (1950).

***Pterulicium sulcisporum*** (Corner) Leal-Dutra, Dentinger, G.W. Griff., **comb. nov**.

MycoBank MB831043

Basionym: *Deflexula sulcispora* Corner, *Ann*.

*Bot*., Lond., n.s. **16**: 283 (1952).

Description in Corner (1952a).

***Pterulicium tenuissimum*** (M.A. Curtis) Leal-Dutra, Dentinger, G.W. Griff., **comb. nov**.

MycoBank MB831044

Basionym: *Typhula tenuissima* M.A. Curtis, *Am. Journ. Art. Scienc*. **6**: 351 (1848).

Synonym: *Pterula tenuissima* (M.A. Curtis) Corner, *Monograph of Clavaria and allied Genera*, (Annals of Botany Memoirs No. 1): 524 (1950).

Description in Corner (1950).

***Pterulicium ulmi*** (Peck) Leal-Dutra, Dentinger, G.W. Griff., **comb. nov**.

MycoBank MB831045

Basionym: *Mucronella ulmi* Peck, *Ann. Rep. Reg. N*.*Y. St. Mus*. **54**: 154 (1902) [1901].

Synonym: *Deflexula ulmi* (Peck) Corner, *Monograph of Clavaria and allied Genera*, (Annals of Botany Memoirs No. 1): 400 (1950).

Description in Corner (1950, 1970).

***Pterulicium velutipes*** (Corner) Leal-Dutra, Dentinger, G.W. Griff., **comb. nov**.

MycoBank MB831046

Basionym: *Pterula velutipes* Corner, *Ann. Bot*., Lond., n.s. **16**: 569 (1952).

Description in Corner (1952b).

