## Supplementary material for "Reclassification of Pterulaceae Corner (Basidiomycota: Agaricales) introducing the ant-associated genus *Myrmecopterula* gen. nov., *Phaeopterula* Henn. and the corticioid Radulomycetaceae fam. nov.": Suppdata2

**Detailed phylogenetic analyses relating to Figure 3**

**IQ-TREE Analysis**

1) IQ-TREE best scheme and substitution models searches, followed by best tree search and UFBoot with 1000 replicates.

Command:

iqtree -s $ALIGNMENT -spp partitions.txt -m MFP+MERGE -bb 1000 -bspec GENESITE

Best_scheme result:

ITS1_ITS2: TPM2u+F+I+G4

58S_LSU: GTR+F+I+G4

RPB2: TIM3e+R3

2) Best scheme and substitution model search to implement on MrBayes

Command:

iqtree -s $ALIGNMENT -spp partitions.txt -m TESTMERGEONLY -mset mrbayes

Best_scheme result:

ITS1_ITS2: GTR+F+I+G4

58S_LSU: GTR+F+I+G4

RPB2: SYM+I+G4

**MrBayes Analysis**

For MrBayes analysis the following files were created. The parameters in the partitions file were retrieved from the IQ-TREE analysis 2 above.

Batch file:

```
begin mrbayes;
   set autoclose=yes nowarn=yes usebeagle=yes beagledevice=GPU beagleprecision=single beaglesse=yes beaglescaling=dynamic beaglethreads=yes;
   execute partitions.nexus;
   mcmc nruns=2 ngen=10000000 nchains=4 burninfrac=0.25 samplefreq=1000 printfreq=1000 diagnfreq=10000 filename=mb_result;
	sump;
	sumt;

end;
```

Partitions file:

```
#nexus
begin mrbayes;
  execute Pter_reduced_19-03_namesok.nexus;
  charset ITS1_ITS2 = 1-464 626-1081;
  charset 58S_LSU = 465-625 1082-1992;
  charset RPB2 = 1993-2740;
  partition favored = 3: ITS1_ITS2, 58S_LSU, RPB2;
  set partition = favored;

  lset applyto=(1,2,3) nst=6 rates=invgamma ngammacat=4;

  prset applyto=(1)
  statefreqpr=fixed(0.226885,0.218572,0.20682,0.347724)
  pinvar=fixed(0.08113342675)
  shapepr=fixed(1.057220686)
  revmat=fixed(1.358794076,3.444954973,1.609693478,0.6406387856,3.853041436,1);

  prset applyto=(2)
  statefreqpr=fixed(0.266692,0.196548,0.287038,0.249722)
  pinvar=fixed(0.5775252228)
  shapepr=fixed(0.5980550403)
  revmat=fixed(1.147009825,4.486162813,2.412293815,0.3723987737,11.19088756,1);

  prset applyto=(3)
  statefreqpr=fixed(equal)
  pinvar=fixed(0.434542201)
  shapepr=fixed(1.383750766)
  revmat=fixed(1.578335371,4.844871367,0.4833771497,1.372958564,7.149962786,1);

end;
```

**Supplementary analysis**

A dataset including all the sequences in Suppdata1 was created to find the best outgroup for Pterulaceae/Radulomycetaceae and to ensure that all the sequences form each cloned samples would cluster together.

The alignment and ML tree reconstruction followed the same methods for the main Figure 3. And the best scheme result was:

ITS1_58S_ITS2_LSU_RPB2: TIM2+F+I+G4

The resulting tree (Suppdata2 Fig. 1) showed Sptephanosporaceae as the best outgroup for Pterulaceae/Radulomycetaceae and

**Suppdata2 Fig. 1 (below):** ML tree of Pterulaceae/Radulomycetaceae and several outgoups.

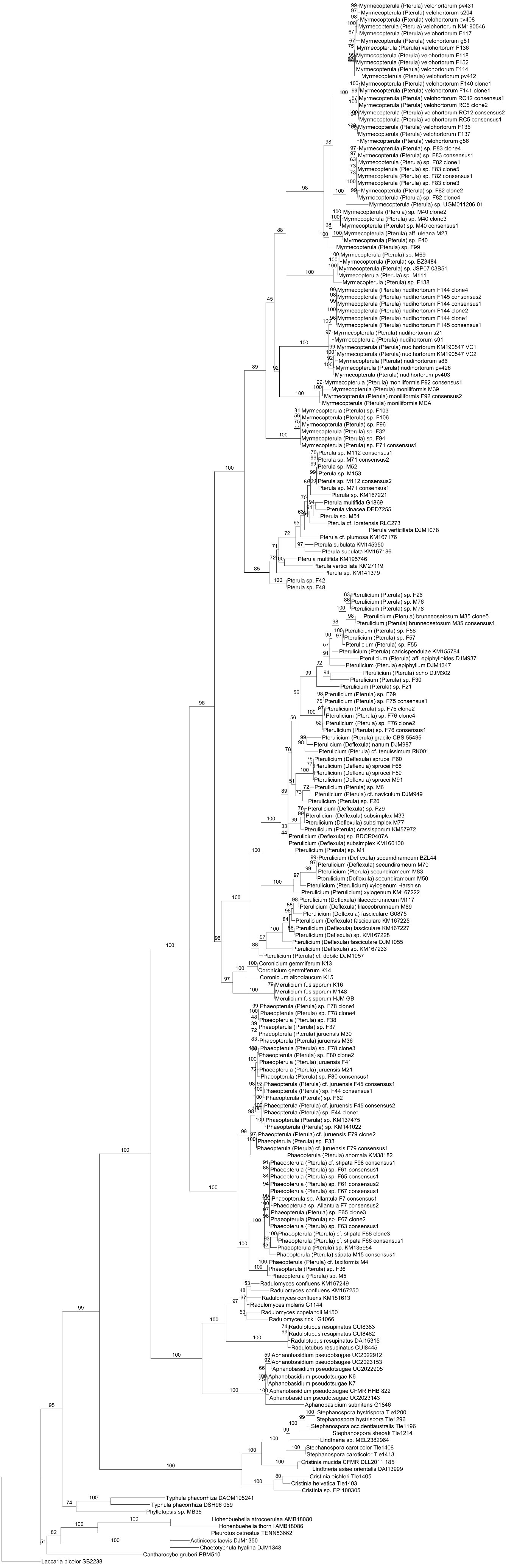

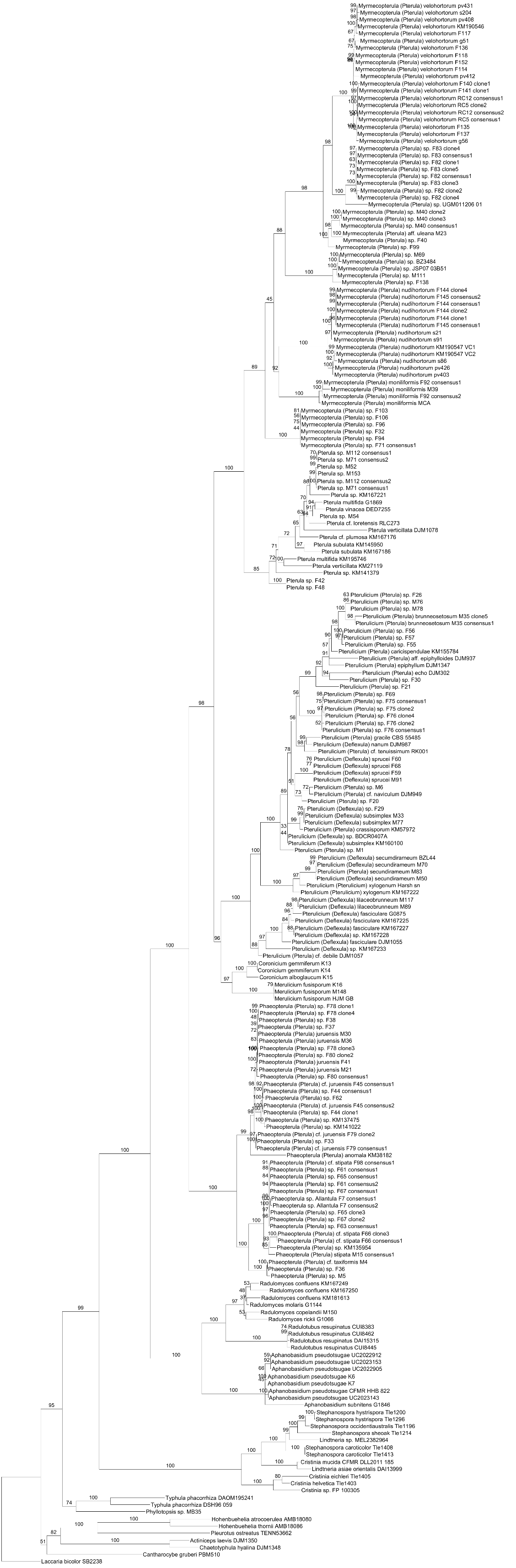
