## Supplementary material for "Reclassification of Pterulaceae Corner (Basidiomycota: Agaricales) introducing the ant-associated genus *Myrmecopterula* gen. nov., *Phaeopterula* Henn. and the corticioid Radulomycetaceae fam. nov.": Suppdata3

### Slide 1
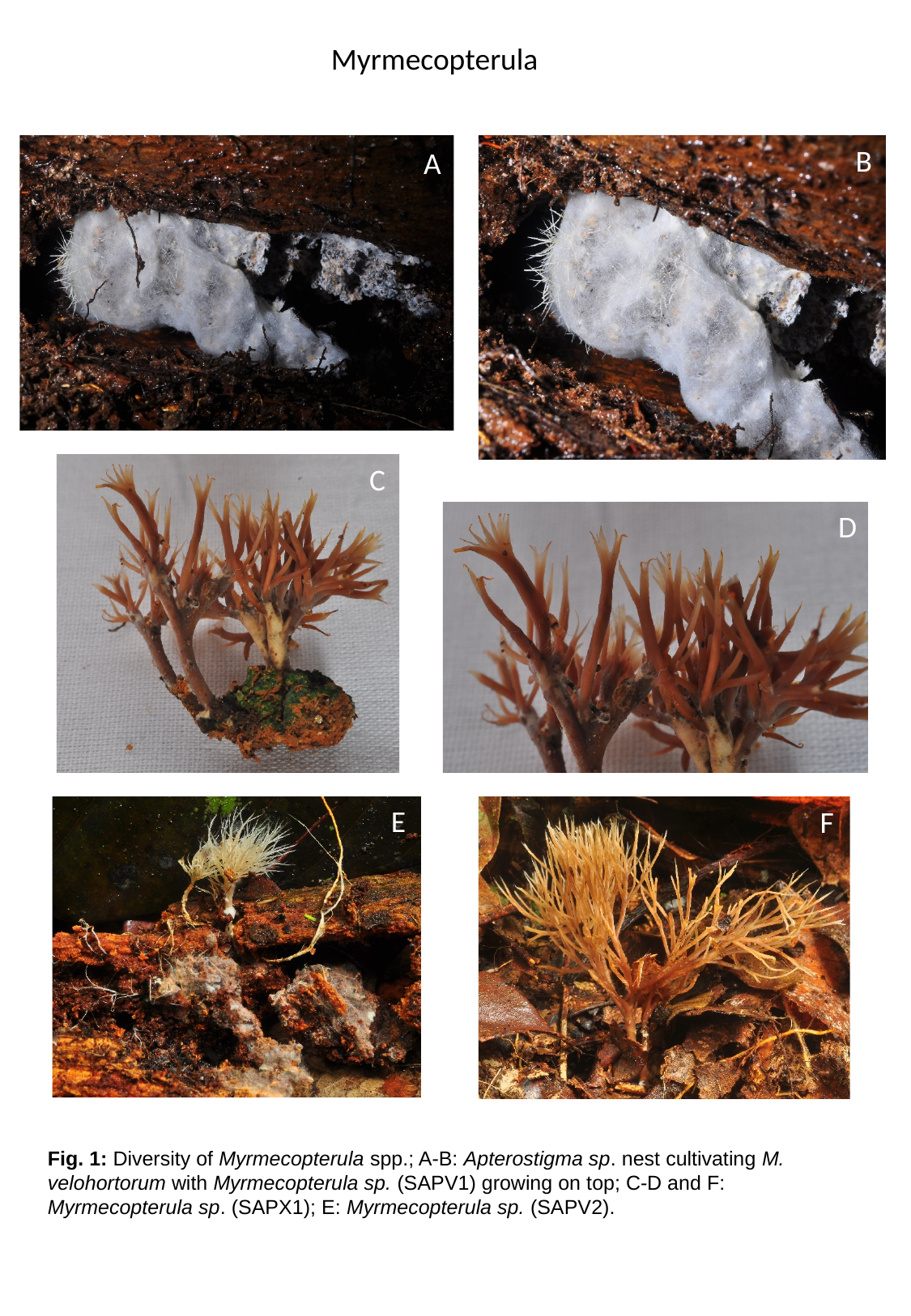

Myrmecopterula
B
A
C
D
E
F
Fig. 1: Diversity of Myrmecopterula spp.; A-B: Apterostigma sp. nest cultivating M. velohortorum with Myrmecopterula sp. (SAPV1) growing on top; C-D and F: Myrmecopterula sp. (SAPX1); E: Myrmecopterula sp. (SAPV2).

### Slide 2
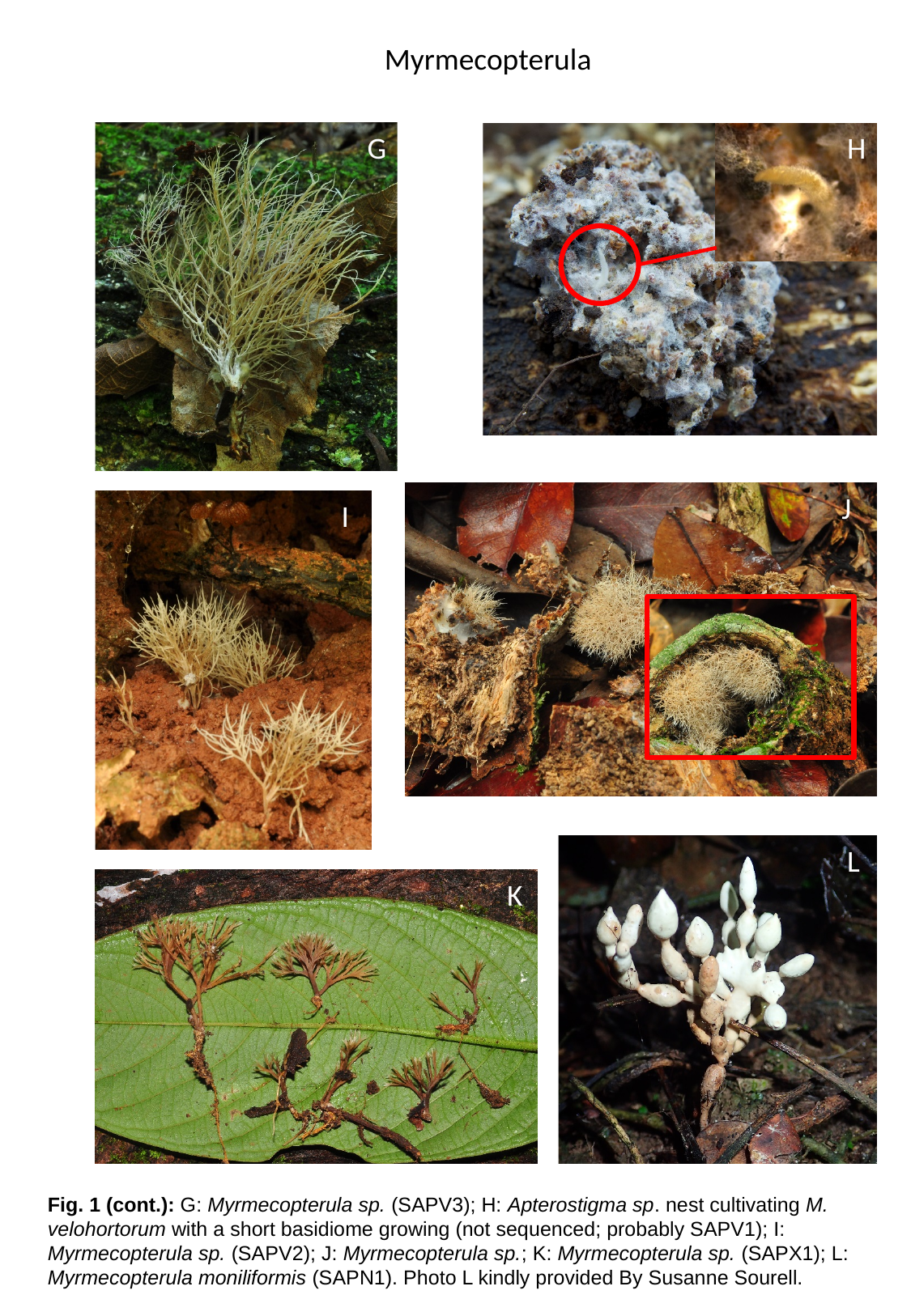

Myrmecopterula
G
H
J
I
L
K
Fig. 1 (cont.): G: Myrmecopterula sp. (SAPV3); H: Apterostigma sp. nest cultivating M. velohortorum with a short basidiome growing (not sequenced; probably SAPV1); I: Myrmecopterula sp. (SAPV2); J: Myrmecopterula sp.; K: Myrmecopterula sp. (SAPX1); L: Myrmecopterula moniliformis (SAPN1). Photo L kindly provided By Susanne Sourell.

### Slide 3
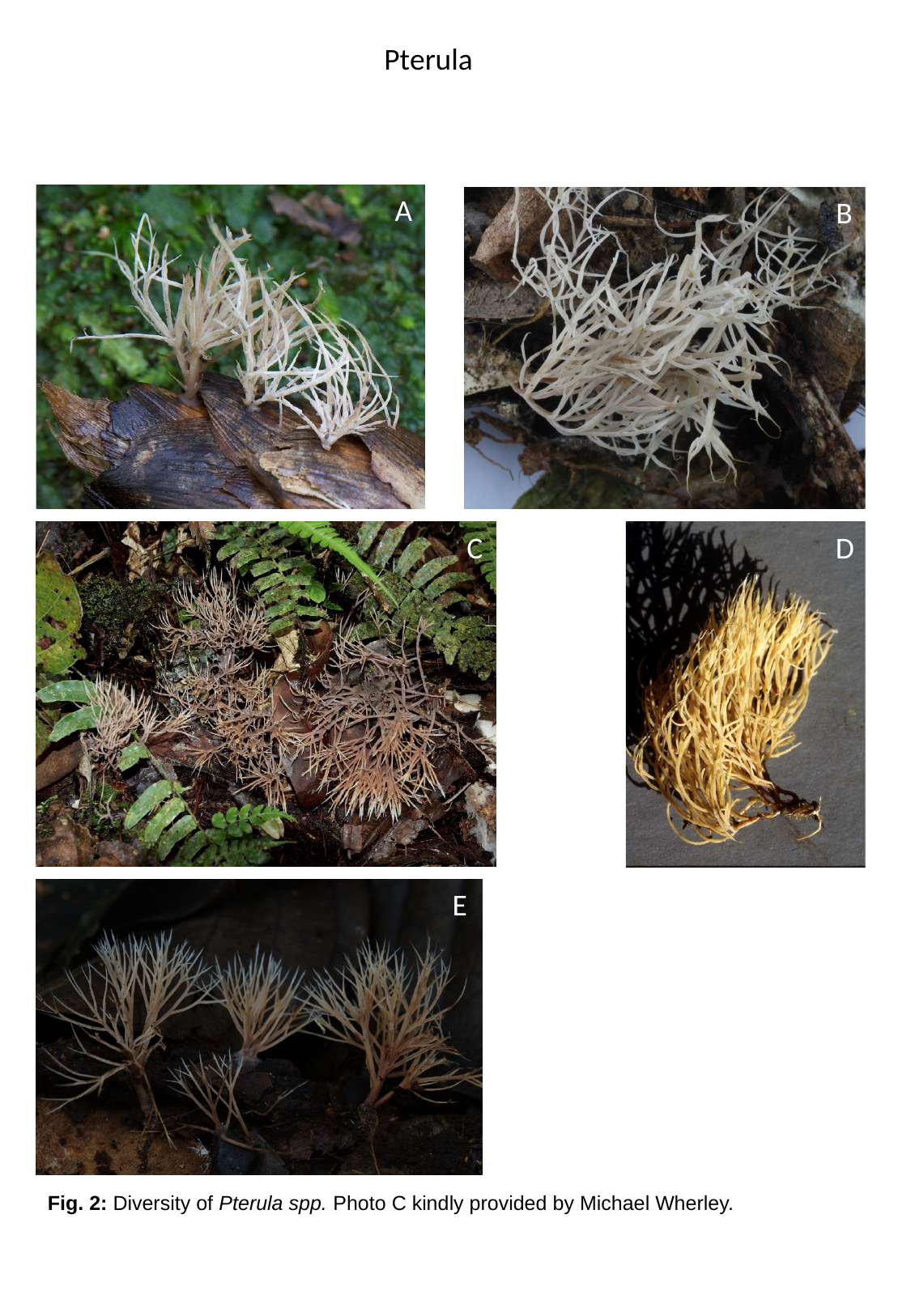

Pterula
A
B
C
D
E
Fig. 2: Diversity of Pterula spp. Photo C kindly provided by Michael Wherley.

### Slide 4
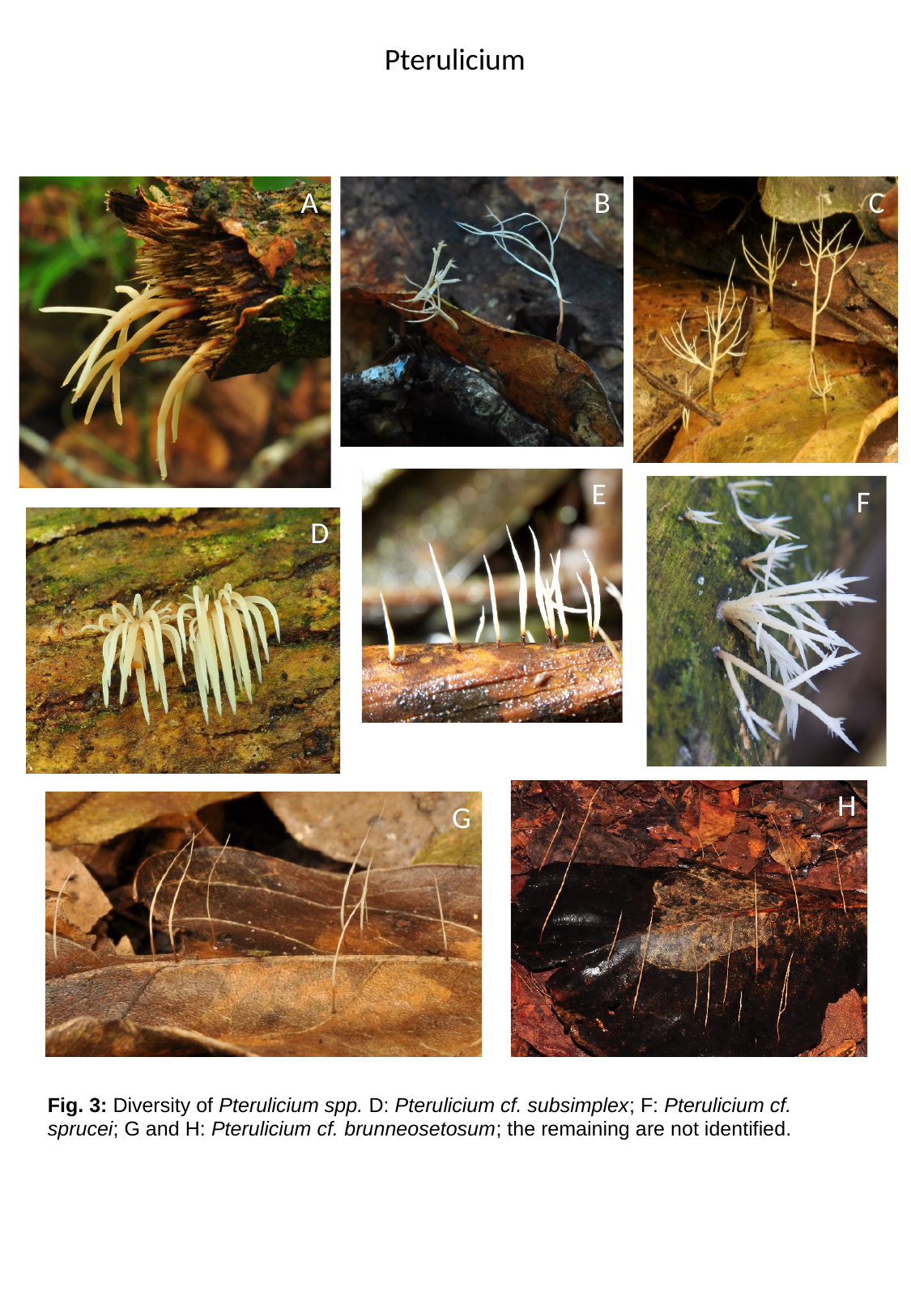

Pterulicium
A
B
C
E
F
D
H
G
Fig. 3: Diversity of Pterulicium spp. D: Pterulicium cf. subsimplex; F: Pterulicium cf. sprucei; G and H: Pterulicium cf. brunneosetosum; the remaining are not identified.

### Slide 5
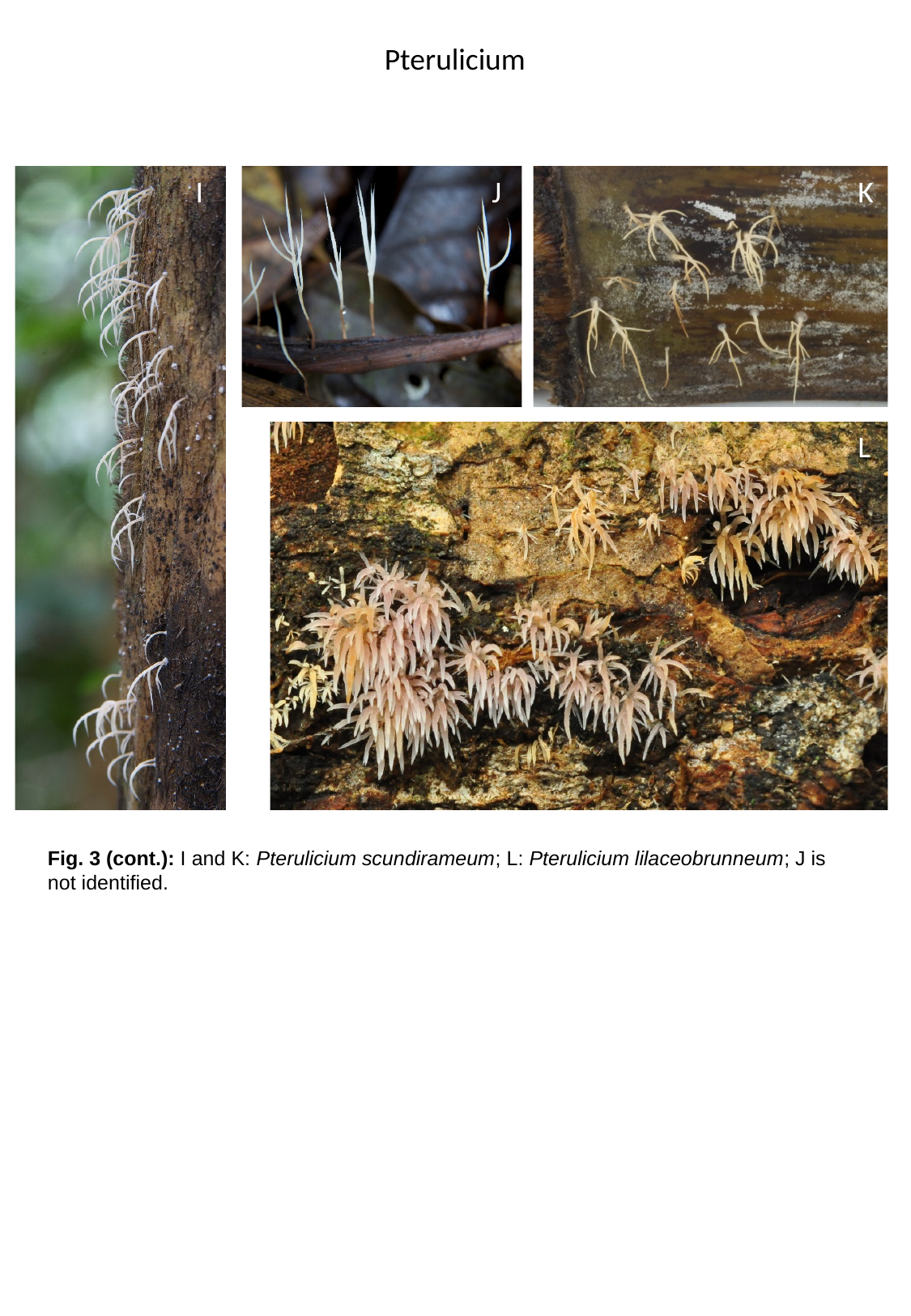

Pterulicium
I
J
K
L
Fig. 3 (cont.): I and K: Pterulicium scundirameum; L: Pterulicium lilaceobrunneum; J is not identified.

### Slide 6
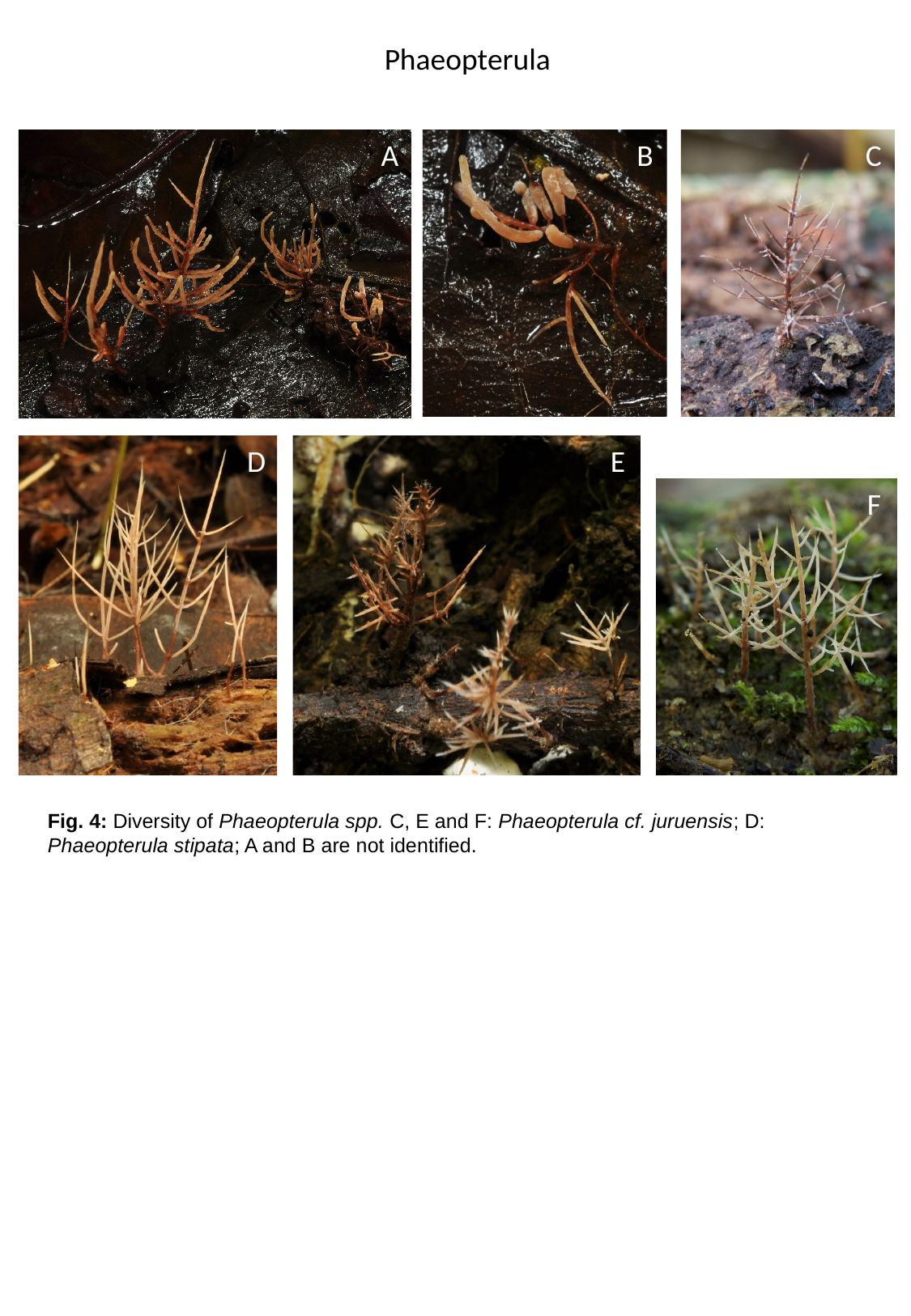

Phaeopterula
A
B
C
D
E
F
Fig. 4: Diversity of Phaeopterula spp. C, E and F: Phaeopterula cf. juruensis; D: Phaeopterula stipata; A and B are not identified.

### Slide 7
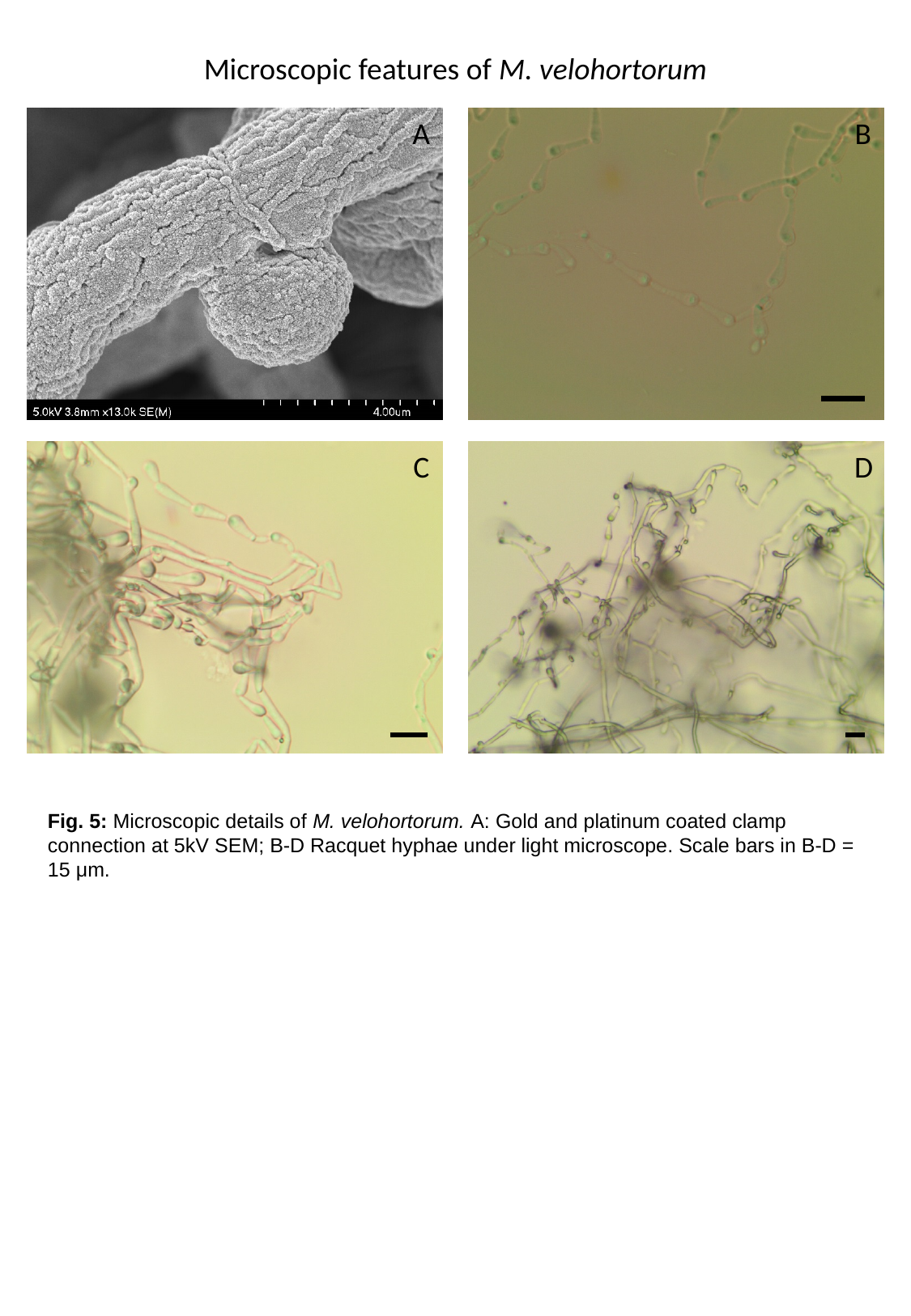

Microscopic features of M. velohortorum
A
B
C
D
Fig. 5: Microscopic details of M. velohortorum. A: Gold and platinum coated clamp connection at 5kV SEM; B-D Racquet hyphae under light microscope. Scale bars in B-D = 15 μm.
