## Supplementary material for "Reclassification of Pterulaceae Corner (Basidiomycota: Agaricales) introducing the ant-associated genus *Myrmecopterula* gen. nov., *Phaeopterula* Henn. and the corticioid Radulomycetaceae fam. nov.": Suppdata4

### ONLINE RESOURCE 4

### ADDITIONAL NOMENCLATURAL NOVELTIES

#### 1) *Phaeopterula*

***Phaeopterula anomala* (P. Roberts) Leal-Dutra, Dentinger, G.W. Griff., comb. nov.**
MycoBank No: 830999

Basionym
*Pterula anomala* P. Roberts, Kew Bull. 54(3): 528 (1999).

Description in Roberts (1999).

***Phaeopterula hirsuta* (Henn.) Sacc. & D. Sacc.**
MycoBank No: 469044

Basionym
*Pterula hirsuta* Henn., in Warburg, Monsunia 1: 9 (1899) [1900]. ≡ *Dendrocladium hirsutum* (Henn.) Lloyd, Mycol. Writ. 5: 870 (1919).

Description in Corner (1950).

***Phaeopterula juruensis* (Henn.) Sacc. & D. Sacc. (Fig. 1O)**
MycoBank No: 634235

Basionym
*Pterula juruensis* Henn. [as '*Phaeopterula juruensis*'], Hedwigia 43(3): 175 (1904). ≡ *Dendrocladium juruense* (Henn.) Lloyd, Mycol. Writ. 5: 870 (1919).

Descriptions in Corner (1950, 1952b).

***Phaeopterula stipata* (Corner) Leal-Dutra, Dentinger, G.W. Griff., comb. nov. (Fig. 1N)**
MycoBank No: 831000

Basionym
*Pterula stipata* Corner, Ann. Bot., Lond., n.s. 16: 568 (1952).

Description in Corner (1952b).

***Phaeopterula taxiformis* (Mont.) Leal-Dutra, Dentinger, G.W. Griff., comb. nov.**
MycoBank No: 831001

Basionym
*Pterula taxiformis* Mont., Syll. gen. sp. crypt. (Paris): 181 (1856). ≡ *Lachnocladium taxiforme* (Mont.) Sacc., Syll. fung. (Abellini) 6: 740 (1888).
= *Pterula humilis* Speg., Revista Argent. Hist. Nat. 1(2): 110 (1891).
= *Pterula humilis* var. *tucumanensis* Speg., Anal. Mus. nac. B. Aires, Ser. 3 12: 280 (1909).

Descriptions in Corner (1950, 1952b).

***Phaeopterula taxiformis* var. *gracilis* (Corner) Leal-Dutra, Dentinger, G.W. Griff., comb. nov.**
MycoBank No: 831002

Basionym
*Pterula taxiformis* var. *gracilis* Corner, Ann. Bot., Lond., n.s. 16: 568 (1952).

Description in Corner (1952b).

#### 2) *Pterulicium*

***Pterulicium argentinum* (Speg.) Leal-Dutra, Dentinger, G.W. Griff., comb. nov.**

MycoBank No: 831003

Basionym
*Mucronella argentina* Speg., Anal. Mus. nac. Hist. nat. B. Aires 6: 178 (1898) [1899]. ≡ *Deflexula argentina* (Speg.) Corner, Ann. Bot., Lond., n.s. 16: 276 (1952).
= *Deflexula lilaceobrunnea* var. *elongata* Corner, Ann. Bot., Lond., n.s. 16: 276 (1952).

Description in (Corner 1950).

***Pterulicium fasciculare*** **(Bres. & Pat.) Leal-Dutra, Dentinger, G.W. Griff., comb. nov.**
MycoBank No: 831016

Basionym
*Pterula fascicularis* Bres. & Pat., Mycol. Writ. 1: 50 (1901). ≡ *Deflexula fascicularis* (Bres. & Pat.) Corner, Monograph of *Clavaria* and allied Genera, (Annals of Botany Memoirs No. 1): 395 (1950).

Description in Corner (1950).

***Pterulicium fluminense*** **(Corner) Leal-Dutra, Dentinger, G.W. Griff., comb. nov.**
MycoBank No: 831017

Basionym
*Pterula fluminensis* Corner, Ann. Bot., Lond., n.s. 16: 567 (1952).

Descriptions in Corner (1952b, 1970).

***Pterulicium gordium*** **(Speg.) Leal-Dutra, Dentinger, G.W. Griff., comb. nov.**
MycoBank No: 831018

Basionym
*Clavaria gordius* Speg., Anal. Soc. cient. argent. 17(2): 83 (1884). ≡ *Pterula gordius* (Speg.) Corner, Monograph of *Clavaria* and allied Genera, (Annals of Botany Memoirs No. 1): 513 (1950).

***Pterulicium oryzae*** **(Remsberg) Leal-Dutra, Dentinger, G.W. Griff., comb. nov.**
MycoBank No: 831033

Basionym
*Pistillaria oryzae* Remsberg, Mycologia 32(5): 668 (1940). ≡ *Pterula oryzae* (Remsberg) Corner, Monograph of *Clavaria* and allied Genera, (Annals of Botany Memoirs No. 1): 519 (1950).

Descriptions in (Corner 1950) and Remsberg (1940)

***Pterulicium phyllodicola*** **(Corner) Leal-Dutra, Dentinger, G.W. Griff., comb. nov.**
MycoBank No: 831034

Basionym
*Pterula phyllodicola* Corner, Beih. Nova Hedwigia 33: 220 (1970).

Description in Corner (1970)

***Pterulicium phyllophilum*** **(McAlpine) Leal-Dutra, Dentinger, G.W. Griff., comb. nov.**
MycoBank No: 831035

Basionym
*Clavaria phyllophila* McAlpine, Agric. Gaz. N.S.W., Sydney 7: 86 (1896). ≡ *Pterula phyllophila* (McAlpine) Corner, Monograph of *Clavaria* and allied Genera, (Annals of Botany Memoirs No. 1): 520 (1950).

***Pterulicium tenuissimum*** **(M.A. Curtis) Leal-Dutra, Dentinger, G.W. Griff., comb. nov.**
MycoBank No: 831044

Basionym
*Typhula tenuissima* M.A. Curtis, Am. Journ. Art. Scienc. 6: 351 (1848). ≡ *Pterula tenuissima* (M.A. Curtis) Corner, Monograph of *Clavaria* and allied Genera, (Annals of Botany Memoirs No. 1): 524 (1950).

Description in Corner (1950).

***Pterulicium ulmi*** **(Peck) Leal-Dutra, Dentinger, G.W. Griff., comb. nov.**
MycoBank No: 831045

Basionym
*Mucronella ulmi* Peck, Ann. Rep. Reg. N.Y. St. Mus. 54: 154 (1902) [1901]. ≡ *Deflexula ulmi* (Peck) Corner, Monograph of *Clavaria* and allied Genera, (Annals of Botany Memoirs No. 1): 400 (1950).

Description in Corner (1950, 1970).

***Pterulicium velutipes*** **(Corner) Leal-Dutra, Dentinger, G.W. Griff., comb. nov.**
MycoBank No: 831046

Basionym
*Pterula velutipes* Corner, Ann. Bot., Lond., n.s. 16: 569 (1952).

Description in Corner (1952b).
